# Flexible Intentions: An Active Inference Theory

**DOI:** 10.1101/2022.04.08.487597

**Authors:** Matteo Priorelli, Ivilin Peev Stoianov

**Affiliations:** Institute of Cognitive Sciences and Technologies National Research Council of Italy Padova, Italy

**Keywords:** Active Inference, Sensorimotor control, Posterior Parietal Cortex, Intentions, Predictive Coding

## Abstract

We present a normative computational theory of how neural circuitry may support visually-guided goal-directed actions in a dynamic environment. The model builds on Active Inference, in which perception and motor control signals are inferred through dynamic minimization of generalized prediction errors. The Posterior Parietal Cortex (PPC) is proposed to maintain constantly updated expectations, or beliefs over the environmental state, and by manipulating them through flexible intentions it is involved in dynamically generating goal-directed actions. In turn, the Dorsal Visual Stream (DVS) and the proprioceptive pathway implement generative models that translate the high-level belief into sensory-level predictions to infer targets, posture, and motor commands. A proof-of-concept agent embodying visual and proprioceptive sensors and an actuated upper limb was tested on target-reaching tasks. The agent behaved correctly under various conditions, including static and dynamic targets, different sensory feedbacks, sensory precisions, intention gains, and movement policies; limit conditions were individuated, too. Active Inference driven by dynamic and flexible intentions can thus support goal-directed behavior in constantly changing environments, and the PPC putatively hosts its core intention mechanism. More broadly, the study provides a normative basis for research on goal-directed behavior in end-to-end settings and further advances mechanistic theories of active biological systems.

## 1 Introduction

Traditional views of sensorimotor control in goal-directed actions like object-reaching involve several steps, starting with target perception, movement planning in the body posture domain, translation of this plan in muscle commands, and only then resulting in movement execution [1]. Predictive Coding views consider prior knowledge of environmental and bodily contexts as providing crucial anticipatory information [2]. Under this perspective, motor control can thus be thought of as beginning with target anticipation, even before obtaining the corresponding sensory evidence. More generally, action could be seen as a continuous process in which motor plans are computed using prior world knowledge and are continuously updated along with the influx of sensory evidence about the body’s state and possible targets.

Intentions encode motor goals - or plans - set before the beginning of motor acts themselves and could be therefore viewed as memory holders of voluntary actions [3, 4, 5, 6]. Several cortical areas handle different aspects of this process: the Premotor cortex (PM) encodes structuring while the Supplementary Motor Area (SMA) controls phasing [7]. In turn, the PPC plays a role in building motor plans and their dynamic tuning, as different PPC neurons are sensitive to different intentions [8]. Notably, intention neurons respond not only when performing a given action but also during its observation, allowing observers to *predict the goal of the observed action and, thus, to “read” the intention of the acting individual* [5]. Motor goals have also been observed down the motor hierarchy, which is an expression of Hierarchical Predictive Coding in the motor domain [9].

In primates, the dorsomedial visual stream provides critical support for continuously monitoring the body posture and the spatial location of objects to specify and guide actions, and for performing visuomotor transformations in the course of evolving movement [10, 11, 12]. The PPC, located at the apex of the dorsal stream, is also bidirectionally connected to frontal areas, motor and somatosensory cortex, placing it in a privileged position to set goal-directed actions and continuously adjust motor plans by tracking moving targets and posture [3, 13] in a common reference frame [14]. Undoubtedly, the PPC plays a crucial role in visually-guided motor control [15, 13, 16] and its more specific subregion V6A is involved in the control of reach-to-grasp actions [17], but its peculiar role is still disputed. The most consistent view is that the PPC estimates the state of the body and the environment and optimizes the body-environment interactions [18]. Others see the PPC as a task estimator [19] or as being involved in endogenous attention and task setting [20]. Its underlying computational mechanism is not fully understood, especially as regards the definition of goals in motor planning and their integration within the control process [21]. For example, the prevailing Optimal Feedback Control theory defines motor goals through task-specific cost functions [22]. Neural-level details of motor-goal coding are becoming increasingly important in light of the growing demand for neural interfaces that provide information about motor intents [7] in support of intelligent assistive devices [23, 24].

To investigate how neural circuitry in the PPC supports sensory guided actions through motor intentions from a computational point of view, we adopted the Active Inference theory of cognitive and motor control, which provides fundamental insights of increasing appeal about the computational role and principles of the nervous system, especially about the perception-action loop [25, 26, 27, 28]. Indeed, Active Inference provides a formalization of these two cortical tasks, both of which are viewed as aiming to resolve the critical goal of all organisms: to survive in uncertain environments by operating within preferred states (e.g., maintaining a constant temperature). Accordingly, both tasks are implemented by dynamic minimization of a quantity called *free energy*, whose process generally corresponds to the minimization of high- and low-level prediction errors, that is, the satisfaction of prior and sensory expectations. There are two branches of Active Inference appropriate to tackle two different levels of control. The discrete framework can explain high-level cognitive control processes such as planning and decision-making - it evaluates expected outcomes to select actions in discrete entities [29]. In turn, dynamic adjustment of action plans in the PPC matches by functionality the Active Inference framework in continuous state space [9, 30]. In short, this theory departs from classical views of perception, motor planning [1], and motor control [22], unifying and considering them as a dynamic probabilistic inference problem [31, 32, 33, 34]. The biologically implausible cost functions typical of Optimal Control theories are replaced by high-level priors defined in the extrinsic state space, allowing complex movements such as walking or handwriting [35, 36].

In the following, we first outline the background computational framework and then elaborate on movement planning and intentionality in continuous Active Inference. Our most critical contributions regard the formalization of goal-directed behavior and the processes linking flexible goals (e.g., dynamic visual targets) with motor plans through the definition of intention functions. We also investigate a more parsimonious approach to motor control based solely on proprioceptive predictions. We then provide implementation details and a practical demonstration of the theoretical contribution in terms of a simulated Active Inference agent, which we show is capable of detecting and reaching static visual goals and tracking moving targets. We also provide detailed performance statistics and investigate the effects of system parameters whose balance is critical to movement stability. Additionally, gradient analysis provides crucial insights into the causes of the movements performed. Finally, we discuss how intentions could be selected to perform a series of goal-directed steps, e.g., a multi-phase action, and illustrate conditions for neurological disorders.

## 2 Computational Background

We first outline the computational principles of the underlying probabilistic and Predictive Coding approach and provide background on variational inference, free energy minimization, Active Inference, and variational autoencoders necessary to comprehend the following main contribution.

### 2.1 The Bayesian Brain Hypothesis

An interesting visual phenomenon, called *binocular rivalry*, happens when two different images are presented simultaneously to each eye: the perception does not conform to the visual input but alternates between the two images. How and why does this happen? It is well-known that priors play a fundamental role in driving the dynamics of perceptual experience, but dominant views of the brain as a feature detector that passively receives sensory signals and computes motor commands have so far failed to explain how such illusions could arise.

In recent years, there has been increasing attention to a radically new theory of the mind called the *Bayesian brain*, according to which our brain is a sophisticated machine that constantly makes use of Bayesian reasoning to capture causal relationships in the world and deliver optimal behavior in an uncertain environment [37, 38, 39]. At the core of the theory is the Bayes theorem, whose application here implies that posterior beliefs about the world are updated according to the product of prior beliefs and the likelihood of observing sensory input. In this view, perception is more than a simple bottom-up feedforward mechanism that detects features and objects from the current sensorium; rather, it comprises a predictive top-down generative model which continuously anticipates the sensory input to test hypotheses and explain away ambiguities.

According to the Bayesian brain hypothesis, this complex task is accomplished by *Predictive Coding*, implemented through message passing of top-down predictions and bottom-up prediction errors between adjacent cortical layers [2]. The former are generated from latent states maintained at the highest levels, representing beliefs about the causes of the environment, while the latter are computed by comparing sensory-level predictions with the actual observations. Each prediction will then act as a cause for the layer below, while the prediction error will convey information to the layer above. It is thanks to this hierarchical organization and through error minimization at every layer that the cortex is able to mimic and capture the inherently hierarchical relationships that model the world. In this view, sensations are only needed in that they provide, through the computation of prediction errors, a measure of how good the model is and a cue to correct future predictions. Thus, ascending projections do not encode the features of a stimulus, but rather how much the brain is *surprised* about it, considering the strict correlation between surprise and model uncertainty.

### 2.2 Variational Bayes

Organisms are supposed to implement model fit or error minimization by some form of variational inference, a broad family of techniques based on the *calculus of variations* and used to approximate intractable posteriors that would otherwise be infeasible to compute analytically or even with classical sampling methods like Monte Carlo [40].

Under the Bayesian brain hypothesis, we can assume that the nervous system maintains latent variables ***z*** about both the unknown state of the external world and the internal state of the organism. By exploiting a prior knowledge *p*(***z***) and the partial evidence *p*(***s***) of the environment provided by its sensors, it can apply Bayesian inference to improve its knowledge [41]. To do so, given the observation **s**, the nervous system needs to evaluate the posterior *p*(***z***|***s***):

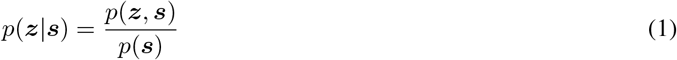

However, directly computing such quantity is infeasible due to the intractability of the marginal *p*(***s***) = *∫ p*(***z, s***)*d**z***, which involves integration over the joint density *p*(***z, s***).

What does the variational approach is approximating the posterior with a simpler to compute *recognition* distribution *q*(***z***) ≈ *p*(***z|s***) through minimization of the Kullback-Leibler (KL) divergence between them:

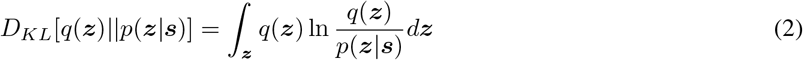

The KL divergence can be rewritten as the difference between log evidence ln *p*(***s***) and a quantity 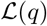 known as *evidence lower bound*, or ELBO [40]:

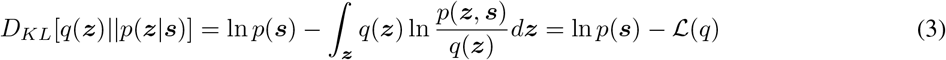

Since the KL divergence is always nonnegative, the ELBO provides a lower bound on log evidence, i.e., 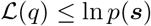. Therefore, minimizing the KL divergence with respect to *q*(***z***) is equivalent to maximizing 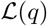, which at its maximum corresponds to an approximate density that is closest the most to the real posterior, depending on the particular choice of the form of *q*(***z***). In general, few assumptions are made about the form of this distribution - a multivariate Gaussian is a typical choice - with a trade-off between having a tractable optimization process and still leading to a good approximate posterior.

### 2.3 Free Energy and Prediction Errors

How can Bayesian inference be implemented through a simple message passing of prediction errors? Friston Friston2002,Friston2005 proposed an elegant solution based on the so-called *free energy*, a concept borrowed from thermodynamics and defined as the negative ELBO. Accordingly, Equation (3) can be rewritten as:

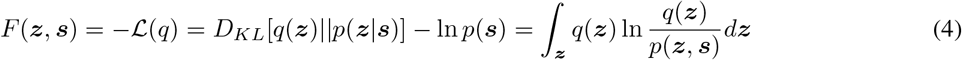

Minimizing the free energy with respect to the latent states ***z*** - a process called *perceptual inference* - is then equivalent to ELBO maximization and provides an upper bound on surprise:

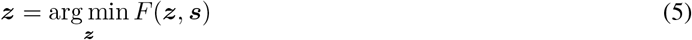

In this way, the organism indirectly minimizes model uncertainty and is able to learn the causal relations between unknown states and sensory input, and to generate predictions based on its current representation of the environment.

Free energy minimization is simpler than dealing with the KL divergence between the approximate and true posteriors as the former depends on quantities the organism has access to, namely the approximate posterior and the *generative model*.

To this concern, it is necessary to distinguish between the latter and the real distribution producing sensory data, called *generative process*, which can be modeled with the following non-linear stochastic equations:

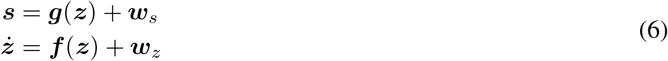

where the function ***g*** maps latent states or causes ***z*** to observed states or sensations ***s***, the function ***f*** encodes the dynamics of the system, i.e., the evolution of ***z*** over time, while ***w**_s_* and ***w**_z_* are noise terms that describe system uncertainty.

Nervous systems approximate the generative process by making a few assumptions: that (i) under the mean-field approximation the recognition density can be partitioned into independent distributions: *q*(***z***) = ⊓*_i_ q*(***z**_i_*), and that (ii) under the Laplace approximation each of these partitions is Gaussian: 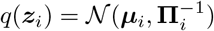, where ***μ**_i_* represents the most plausible hypothesis - also called *belief* about the hidden state ***z**_i_* - and **⊓***_i_* is its precision matrix [42]. In this way, the free energy does not depend on ***z*** and simplifies as follows:

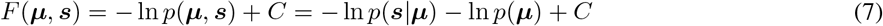

where *C* is a constant term.

A more precise description of the unknown environmental dynamics can be achieved by considering not only the 1st-order of Equation (6) but also higher temporal orders of the corresponding approximations: 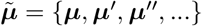 - called *generalized coordinates* [43, 44]. This allows us to better represent the environment with the following generalized model:

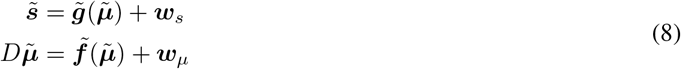

where *D* is the differential (shift) operator matrix such that 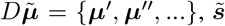 denotes the generalized sensors, while 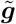 and 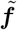 denote the generalized model functions of all temporal orders. Note that in this system, the sensory data at a particular dynamical order ***s***^[*d*]^ - where [*d*] is the order - engage only with the same order of belief ***μ***^[*d*]^, while the generalized equation of motion, or system dynamics, specifies the coupling between adjacent orders. Such equations are generated from the generalized likelihood and prior distributions, which can be expanded as follows:

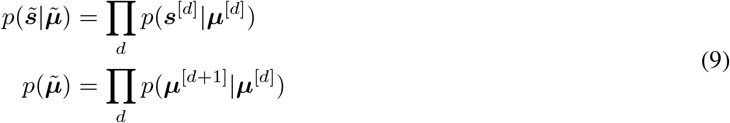

As defined above, these variational probability distributions are assumed to be Gaussian:

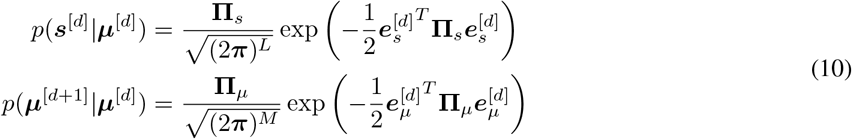

where *L* and *M* are the dimensions of sensations and internal beliefs, respectively with precisions **⊓***_s_* and **⊓***_μ_*. Note that the probability distributions are expressed in terms of sensory and dynamics prediction errors:

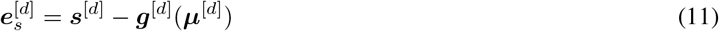

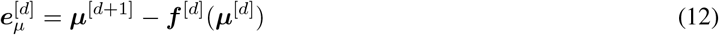

The factorized probabilistic approximation of the dynamic model allows easy state estimation performed by iterative gradient descent over the generalized coordinates, that is, by changing the belief 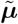 over the hidden states at every temporal order:

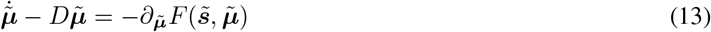

Gradient descent is tractable because the Gaussian variational functions are smooth and differentiable and the derivatives are easily computed in terms of generalized prediction errors, since the logarithm of Equation (7) vanishes the exponent of the Gaussian. The belief update thus turns to:

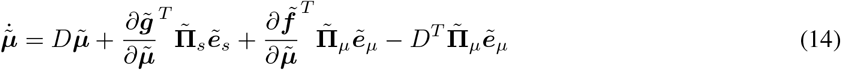

It is crucial to keep in mind the nature of the three components that compose this update equation: a likelihood error computed at the sensory level, a backward error arising from the next temporal order, and a forward error coming from the previous order. These terms represent the free energy gradients relative to the belief ***μ***^[*d*]^ of Equation (11) for the likelihood, and ***μ***^[*d*+1]^ and ***μ***^[*d*]^ of Equation (12) for the dynamics errors.

In short, by making a few plausible simplifying assumptions, the complexity of free energy minimization reduces to the generation of predictions, which are constantly compared with sensory observations to determine a prediction error signal. This error then flows back through the cortical hierarchy to adjust the distribution parameters accordingly and minimize sensory surprise - or maximize evidence - in the long run.

### 2.4 Active Inference

Describing the relationship between Predictive Coding and Bayesian inference still does not explain why the cortex has evolved in such a peculiar way. The answer comes from the so-called *free energy principle* (FEP), regard to which the Bayesian brain hypothesis is just a corollary. Indeed, learning the causal relationships of some observed data (e.g., what causes an increase in body temperature) is insufficient to keep organisms alive (e.g., maintaining the temperature in a vital range).

The FEP states that, for an organism to maintain a state of homeostasis and survive, it must constantly and actively restrict the set of latent states in which it lives to a narrow range of life-compatible possibilities, counteracting the natural tendency for disorder [45] - hence the relationship with thermodynamics. If these states are defined by the organism’s phenotype, from the point of view of its internal model they are exactly the states that it expects to be less surprising. Thus, while perceptual inference tries to optimize the belief about hidden causes to explain away sensations, if on the other hand the assumptions defined by the phenotype are considered to be the true causes of the world, interacting with the external environment means that the agent will try to sample those sensations that make the assumptions true, fulfilling its needs and beliefs. *Active inference* becomes a self-fulfilling prophecy. In this view, there is no difference between a desire and a belief: we simply seek the states in which we expect to find ourselves [26, 46].

For achieving a goal-directed behavior, it is then sufficient to apply free energy minimization also with respect to the action (see Equation 7):

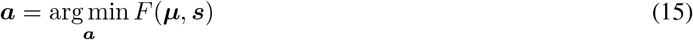

Given that the motor control signals only depend on the sensory information, we obtain:

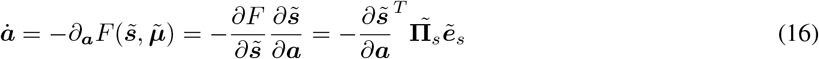

Minimizing the free energy of all sensory signals is certainly useful, as every likelihood contribution will drive the belief update; however, it requires the knowledge of an inverse mapping from exteroceptive sensations to actions [47], which is generally considered a “hard problem” since the mapping is highly non-linear and not univocal [26].

In a more realistic scenario, only proprioception drives the minimization of free energy with respect to the motor signals; this process is easier to realize since the corresponding sensory prediction is already in the intrinsic domain. Control signals sent from the motor cortex are then not motor commands as in classical views of Optimal Control theories; rather, they consist of predictions that define the desired trajectory. Under this perspective, proprioceptive prediction errors computed locally at the spinal cord serve two purposes that only differ in how these signals are conveyed. They drive the current belief toward sensory observations - happening to realize perception - like for exteroceptive signals. But they also drive sensory observations toward the current belief by suppression in simple reflex arcs that activate the corresponding muscles - thus happening to realize movement [36, 48, 49].

In conclusion, perception and action can be seen as two sides of the same coin implementing the common vital goal of minimizing entropy or average surprise. In this view, what we perceive never tries to perfectly match the real state of affairs of the world, but is constantly biased toward our preferred states. This means that action only indirectly fulfills future goals; instead, it continuously tries to fill the gap between sensations and predictions generated from our already biased beliefs.

### 2.5 Variational Autoencoders

Variational Autoencoders (VAEs) belong to the family of generative models, since they learn the joint distribution *p*(***z, s***) and can generate synthetic data similar to the input, given a prior distribution *p*(***z***) over the latent space. VAEs use the variational Bayes approach to capture the posterior distribution *p*(***z|s***) of the latent representation of the inputs when the computation of the marginal is intractable [50]. A VAE is composed of two probability distributions, both of which are assumed to be Gaussian: a probabilistic *encoder* corresponding to the recognition distribution *q*(***z|s***), and a generative function *p*(***s|z***) called probabilistic *decoder* computing a distribution over the input space given a latent representation ***z*** (Figure 3C):

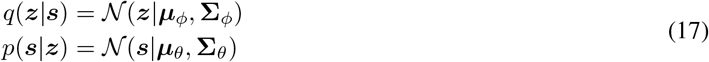

Although VAEs have many similarities with traditional autoencoders, they are actually a derivation of the AEVB algorithm when a neural network is used for the recognition distribution [51].

Unlike other variational techniques, the approximate posterior is generally not assumed to be factorial, but since the calculation of 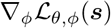 is biased, a method called *reparametrization trick* is used to rewrite the gradient so that it is independent of the parameters ***ϕ***.

This method works by expressing the latent variable ***z*** by a function:

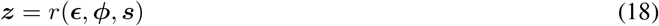

where ***ϵ*** is an auxiliary variable independent of ***ϕ*** and ***s***. The ELBO for a single data point can thus be expressed as:

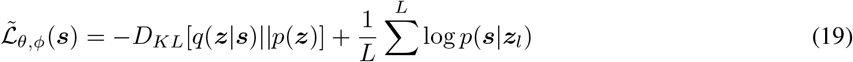

which can be minimized through backpropagation. Here, the KL divergence can be seen as a regularizer, while the second RHS term is an expected negative reconstruction term.

## 3 Results

In what follows, we develop and preliminarily assess a computational theory of the circuitry of the PPC and related areas controlling goal-directed actions in a dynamically changing environment through flexible intentions. We first elaborate on intentionality in Active Inference, then provide a proof-of-concept agent endowed with visual input, and finally describe and analyze several simulations of visually-guided behavior. The theoretical work is motivated by basic research showing the critical role of the PPC in goal-directed sensorimotor control through intention coding [3, 16, 11] and extends previous theoretical and applied research on Active Inference [52, 53, 54] and VAE-based vision support [55, 56, 57]. In turn, the simulations were inspired by a monkey reaching task [58].

### 3.1 Flexible Intentions

State-of-the-art implementations of continuous Active Inference have proven to successfully tackle a wide range of tasks, from oculomotion dynamics [59] to the well-known mountain car problem [52]. Most simulations involve reaching movements in robotic experiments, where several strategies have been tried for designing goal states. These are defined in terms of an attractor embedded in the system dynamics, which can either be a desired latent state [60] or a low-level sensory prediction that is compared with the current observation and backpropagated to obtain a belief update direction [56, 61, 53]. In the simplest case, the goal state is static and the agent is not able to deal with continuously changing environments, expecting that the world will always evolve in the same way. For dynamic goals, one has to use low-level information of sensory signals (e.g., a visual input about a moving target) straight into the high-level dynamics function [35]. When the attractor is in this domain, one has also to duplicate the considered exteroceptive generative model to compare the belief with the dynamic target - which is fine as long as the generative model is fixed, but problems arise when it can be changed by learning.

This is an issue involving biological plausibility: how does dynamic sensory information get available for generating high-level dynamic goals? The same inference process of environmental causes should be at work for the same signal flow, and a goal state should be computed locally without information passed inconsistently. How to design then a dynamic target that avoids implausible scenarios?

Although the high-level latent state could be as simple as encoding body configurations only, an agent could maintain a dynamically estimated belief over moving objects in the scene. An intention can then be computed by exploiting this new information to compute a future action goal in terms of body posture, so that the attractor - either defined in the belief domain or at the sensory level - is not fixed but depends on current perceptual and internal representations of the world - but also on past memorized experiences. The way such an intention is generated may also depend on priors generated from higher-level areas [9]: the considered belief is then located at an intermediate level between the generative models that produce sensory predictions, and the ones that define its evolution over time.

But there is no need to constrain the belief dynamics to a single intention: we propose to decompose it into a set of functions, each one providing an independent expectation that the agent will find itself in a particular state. The belief is then constantly subject to several forces of two natures: one from below - proportional to the *sensory prediction errors* - that pulls it toward what the agent is currently perceiving, and one from higher hierarchical levels - which we call *intention prediction errors* - that pulls it toward what the agent expects to perceive in the future.

As shown in Figure 1, from a neural perspective the PPC is the ideal candidate for a cortical structure computing beliefs over bodily states and flexible intentions: on one hand, being at the apex of the DVS and other sensory generative models, and on the other linked with motor and frontal areas that produce continuous trajectories and plans of discrete action chunks. The PPC is known to be an associative region that integrates information from multiple sensory modalities and encodes visuomotor transformations - e.g., area V6A is thought to encode object affordances during reaching and grasping tasks [62, 10]. Moreover, evidence suggests that the PPC encodes multiple goals in parallel during sequences of actions, even when there is a considerable delay between different goals [63].

**Figure 1:**
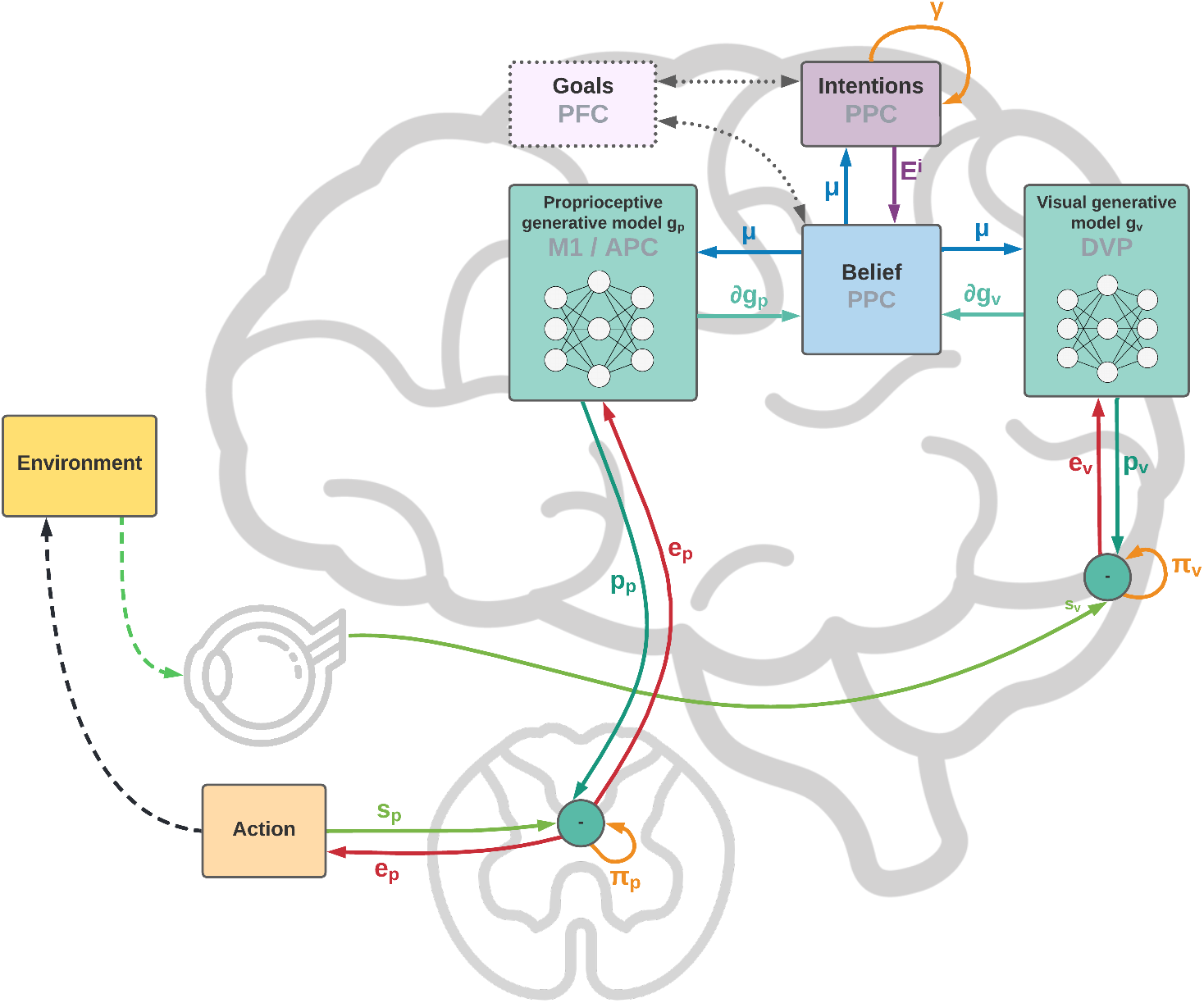
Functional architecture and cortical overlay. The process starts with the computation of future intentions ***h*** in the PPC under the coordination of frontal and motor areas. In the middle of the sensorimotor hierarchy, the PPC maintains beliefs ***μ*** over the latent causes of sensory observations ***s**_p_* and ***s**_v_*, and computes proprioceptive and visual predictions through the somatosensory and dorsal visual pathways (for simplicity, we have omitted the somatomotor pathway and considered a single mechanism for both motor control and belief inference). The lower layers of the hierarchy compute sensory prediction errors ***e**_p_* and ***e**_v_*, while the higher layers compute intention prediction errors ***e***^(*i, k*)^; both are propagated back toward the PPC, which thus integrates information from multiple sensory modalities and intentions. Free energy is minimized throughout the cortical hierarchy by changing the belief about the causes of the current observation (perception) and by sending proprioceptive predictions from the motor cortex to the reflex arcs (action). An essential element of this process is the computation of gradients *∂**g**_p_* and *∂**g**_v_* by inverse mappings from the sensations toward the deepest latent states. In this process, intentions act as high-level attractors and the belief propagated down to compute sensorimotor predictions embeds a component directing the body state toward the goals.

In short, the agent maintains plausible hypotheses over the causes of its percepts, either bodily states or objects in the exteroceptive domains; by manipulating them, the agent dynamically constructs representations of future states, i.e., intentions, which in turn act as priors over the current belief. Thus, if the job of the sensory pathways is to compute sensory-level predictions, we hypothesize that higher levels of the sensorimotor control hierarchy integrate in the PPC previous states of belief with flexible intentions, each predicting the next plausible belief state.

For a more formal definition, we assume that the neural system perceives the environment and receives motor feedback through *J* noisy sensors ***s*** comprising multiple domains (most critically, proprioceptive and visual). Under the VB and Gaussian approximations of the recognition density, we also assume that the nervous system operates on beliefs ***μ*** ∈ ℝ*^M^* that define an abstract internal representation of the world. Furthermore, we assume that the agent maintains generalized coordinates up to the 2nd-order resulting from free energy minimization in the generalized belief space 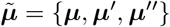.

We then define *intentions **h*** as predictions of target goal states over current beliefs ***μ*** computed with the help of *K intention functions **i***^(*k*)^(***μ***) ∈ ℝ*^M^* building intention representations ***h***^(*k*)^ for each goal of the agent. Although both belief and intentions could be abstract representations of the world - comprising states in extrinsic and intrinsic coordinates - we assume a simpler scenario in which the intentions operate on beliefs in a common intrinsic motor-related domain, e.g., the joint angles space. As explained before, we assume that there are two conceptually different components in both the belief ***μ*** and the output of the intention functions ***i***^(*k*)^. The first component could represent the bodily states and serve to drive actions, while the second one could represent the state of other objects - mostly targets to interact with - which can be internally encoded in the joint angles space as well (the reason for this particular encoding will be clear later). These targets could be observed, but they could also be imagined or set by higher-level cognitive control frontal areas such as the PFC or PMd [64, 65].

For the sake of notational simplicity, we group all intentions into a single matrix ***H*** ∈ ℝ*^M x K^*:

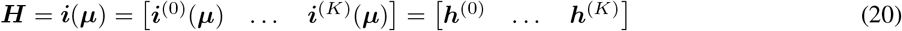

Intention prediction errors ***e***^(*i, k*)^ are then defined as the difference between the current belief and every intention:

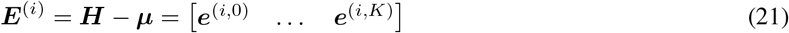

In turn, sensory predictions are produced by a set of generative models ***g**_j_*, one for each sensory modality. We group the predictions into a prediction vector ***p***:

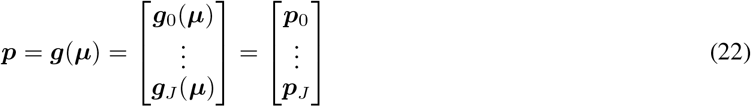

Note that each term ***p**_j_* is a multidimensional sensory-level representation that provides predictions for a particular sensory domain, with its own dimensionality, which we group into a single quantity for notational simplicity.

Sensory prediction errors ***e**_j_* are then computed as the difference between sensations from each domain and the corresponding sensory-level predictions:

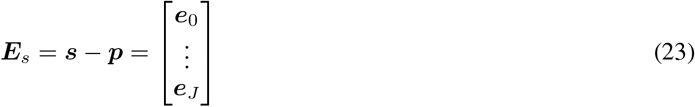

In the above notation and hereafter, the superscripts index intentions, while the subscripts index sensory domains.

### 3.2 Dynamic Goal-directed Behavior in Active Inference

Under the assumption of independence among intentions and sensations, we can factorize the joint probability of the generative model into a product of distributions for each sensory modality and intention, which expands as follows:

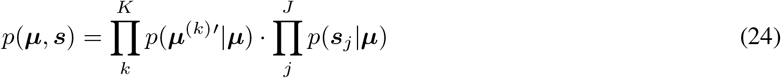

The probability distributions are assumed to be Gaussian:

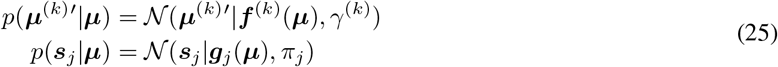

where *γ*^(*k*)^ and *π_j_* are, respectively, the precisions of intention *k* and sensor *j*. Here, ***μ***^(*k*)’^ and ***f***^(*k*)^ correspond to the kth component of the generalized dynamics function:

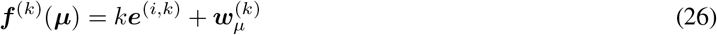

where *k* is the gain of intention prediction errors ***E***^(*i*)^. Note that the goal states are embedded into the 1st-order dynamics functions, acting as belief-level attractors for each intention, so that the agent expects to be pulled toward target states with a velocity proportional to the error.

Although the generalized belief allows encoding information about the dynamics of the true generative process, in the simple case delineated the agent does not have any such prior. For example, the agent does not know the trajectory of a moving target in advance (whose prior, in a more realistic scenario, would be present and acquired through learning of past experiences) and will update the belief only relying on the incoming sensory information. Nevertheless, the agent maintains (false) expectations about target dynamics, and it is indeed the discrepancy between the evolution of the (real) generative process and that of the (internal and biased) generative model that makes it able to implement a goal-directed behavior.

The prediction errors of the dynamics functions can be grouped into a single matrix:

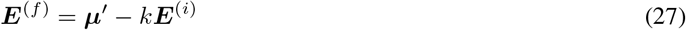

From Equation 14, we can now compute the free energy derivative with respect to the belief:

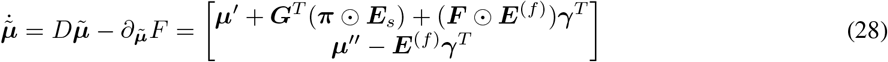

Here, ʘ is the element-wise product, ***G*** and ***F*** enclose the gradients of all sensory generative models and dynamics functions, while ***π*** and ***γ*** comprise all sensory and intention precisions:

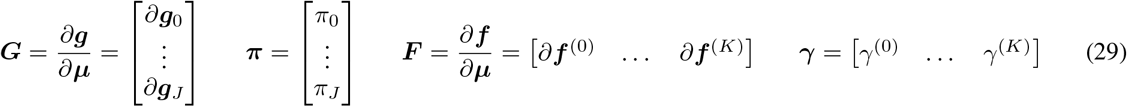

In the following, we will neglect the backward error in the 0th-order of Equation (28) since it has a much smaller impact on the overall dynamics, and treat as the actual attractor force the forward error at the 1st-order:

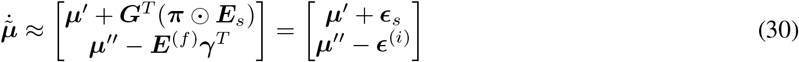

where ***ϵ**_s_* and ***e***^(*i*)^ respectively stand for the contributions of precision-weighted sensory and intention prediction errors. Considering the 1st-order forward error as attractive force instead of the 0th-order backward error - as in most state-of-the-art implementations - also results in computational advantages since there is no gradient of the dynamics functions to be considered. Further studies are however needed to understand the relationships between these two forces in goal-directed behavior.

We can interpret ***γ***^(*k*)^ as a quantity that determines the relative attractor gain of intention *k*, so that intentions with greater strength have a more significant impact on the overall update direction; these gains could also be modulated by projections from higher-levels areas applying cognitive control. In turn, *π_j_* corresponds to the confidence about each sensory modality *j*, so that the agent relies more on sensors with higher strength.

Similarly, we can compute control signals by minimizing the free energy with respect to the actions, expressing the mapping from sensations to actions by:

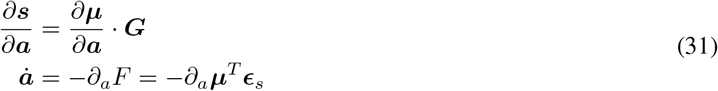

where *∂_a_**μ*** is an inverse model from belief to actions. If motor signals are defined in terms of joint velocities, we can decompose and approximate the inverse model as follows:

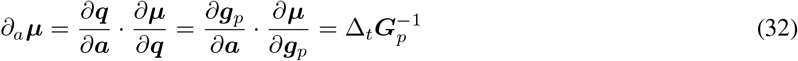

where the subscript p indicates the proprioceptive contribution and we approximated *∂_a_**g**_p_* by a time constant Δ*_t_* [61].

If we assume that the belief over hidden states is encoded in joint angles, the computation of the inverse model may be as simple as finding the pseudoinverse of a matrix. However, if the belief is specified in a more generic reference frame and the proprioceptive generative model is a non-linear function, it could be harder to compute the corresponding gradient, causing additional control problems like temporal delays on sensory signals [35].

Alternatively, we can consider a motor control driven only by proprioceptive predictions, so that the control signal is already in the correct domain and may be achieved through simple reflex arc pathways [36, 49]. In this case, all that is needed is a mapping from proprioceptive predictions to actions:

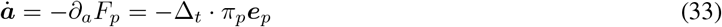

Expressing in Equation (31) the mapping from sensations to actions by the product of the inverse model *∂_a_**μ*** and the gradient of the generative models allows the control signals to be defined in terms of the weighted sensory contribution ***ϵ**_s_,* already computed during the inference process. As will be explained later, such an approach may have some advantages but it is unlikely to be implemented in the nervous system, as control signals do not convey predictions anymore and cannot be realized through reflex arcs at low levels of the motor hierarchy.

Algorithm 1 outlines a schematic description of the flow of dynamic computations. For simplicity, we used the term “intention” also when describing the dynamics functions and their precisions, but one has to keep in mind the difference between the intention prediction errors ***E***^(*i*)^, which directly encode the direction toward target states, and the dynamics prediction errors ***E***^(*f*)^, which arise from the derivation of the corresponding probability distributions.

#### Algorithm 1 Active Inference agent with flexible intentions

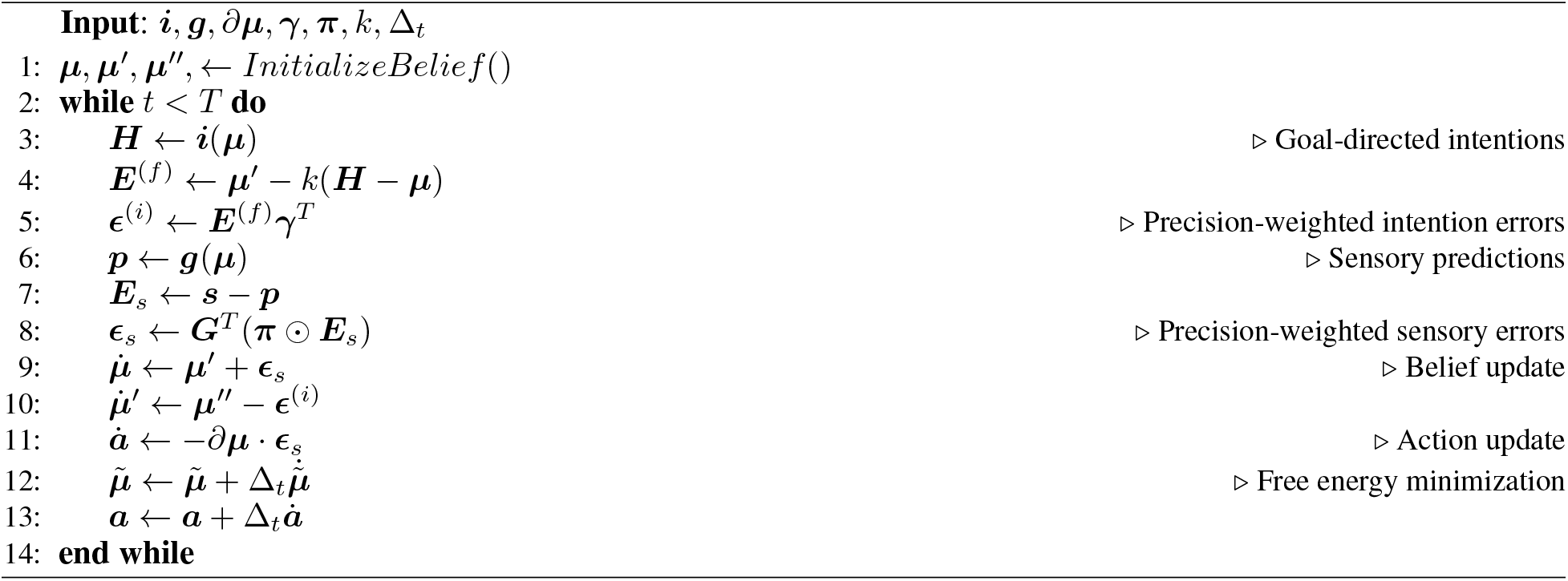

### 3.3 Neural Implementation

Figure 2 shows a schematic neuronal representation of the proposed agent, which further extends earlier perceptual inference schemes [27] to full-blown Active Inference. In this simple model, the intentions consist of a single layer with two neurons, and the goal states are implicitly defined in the dynamics functions; in a realistic setting however the latter would be composed of networks of neurons where these states are explicitly encoded, and non-linear functions could also be used to achieve more advanced behaviors. Note also that intentions ***h***^(*k*)^ and sensory generative models ***g**j* are all part of the same architecture, the only difference being the location in the cortical hierarchy.

**Figure 2:**
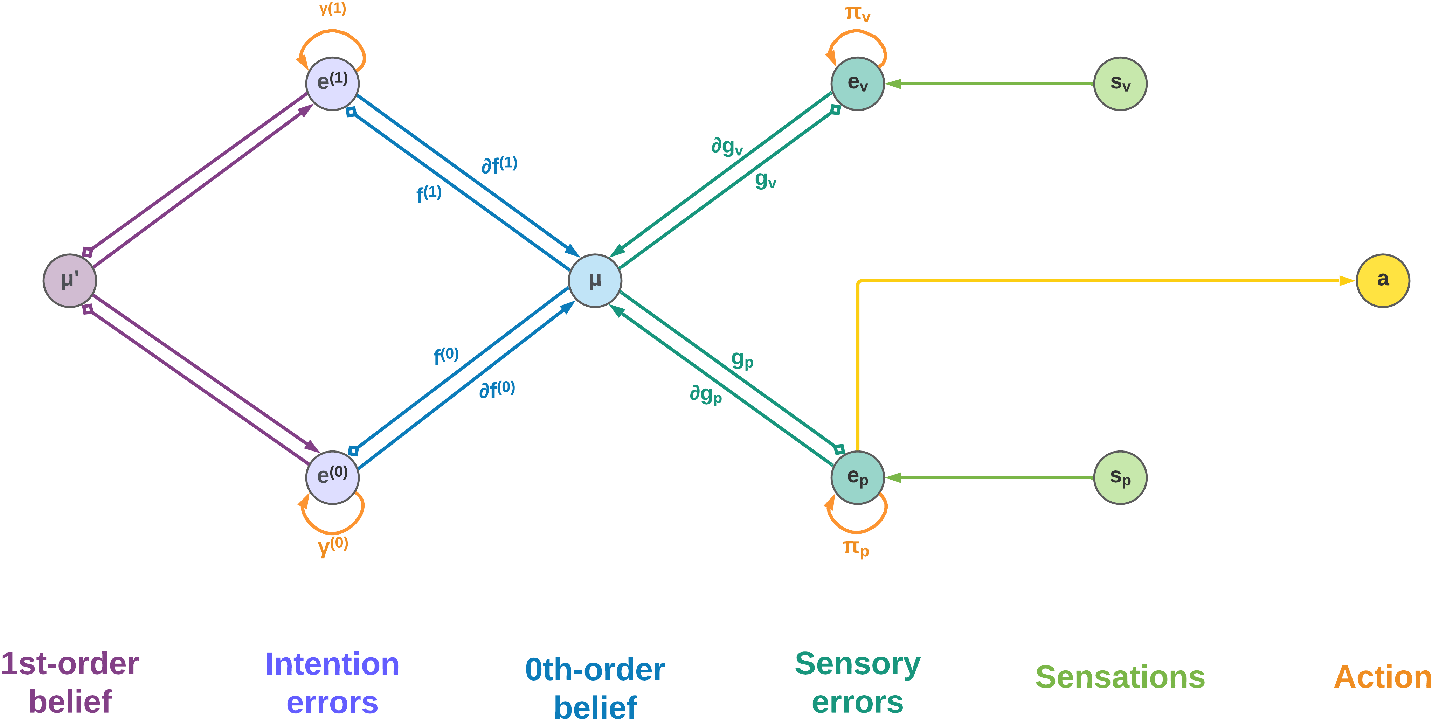
Neuronal representation with two intentions. Small squares stand for inhibitory connections.

Low-level prediction errors for each sensory modality are represented by neurons whose dynamics depends on both observations and predictions of the sensory generative models:

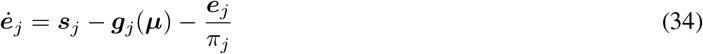

Upon convergence of the neural activity, that is, 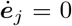, we obtain the prediction error computation derived above.

In turn, the internal activity of neurons corresponding to high-level prediction errors is obtained by subtracting the generated dynamics function from the 1st-order belief:

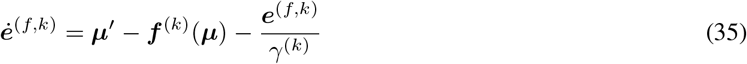

Having received information coming from the top and bottom of the hierarchy, the belief is updated by integrating every signal:

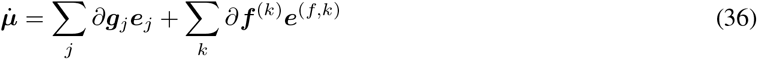

which parallels the update formula derived above (Equation 30).

Correspondingly, the 1st-order component of the belief is updated as follows:

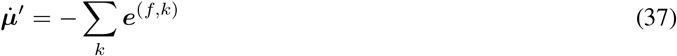

The belief is thus constantly pushed toward a direction that matches sensations on one side and intentions on the other. We adopted the idea that the slow-varying precisions are encoded as synaptic strengths [27], but alternative views consider them as gains of superficial pyramidal neurons [66]. In any case, they can be dynamically optimized during inference in a direction that minimizes free energy - e.g., if a sensory modality does not help predict sensations, its weight will decrease. This is also true for the intention weights: by dynamically changing during the movement, they can act as modulatory signals that select the best intention to realize at every moment (e.g., for solving simultaneous or sequential tasks). Nonetheless, the distinction is purely conceptual as the agent does not discriminate between modulating a future action signal or increasing the confidence of a sensory signal. At the belief level, every element just follows the rules of free energy minimization.

### 3.4 Simulated Agent

To demonstrate the feasibility of the approach, we simulated an agent consisting of an actuated upper limb with visual and proprioceptive sensors that allow it to perceive and reach static and moving targets within its reach. Figure 3A shows the size and position of the targets, as well as limb size and a sample posture. Since the focus here was on theoretical aspects, we simulated just a coarse 3-DoF limb model moving on a 2D plane. However, the approach easily generalizes to a more elaborated limb model and 3D movements.

**Figure 3:**
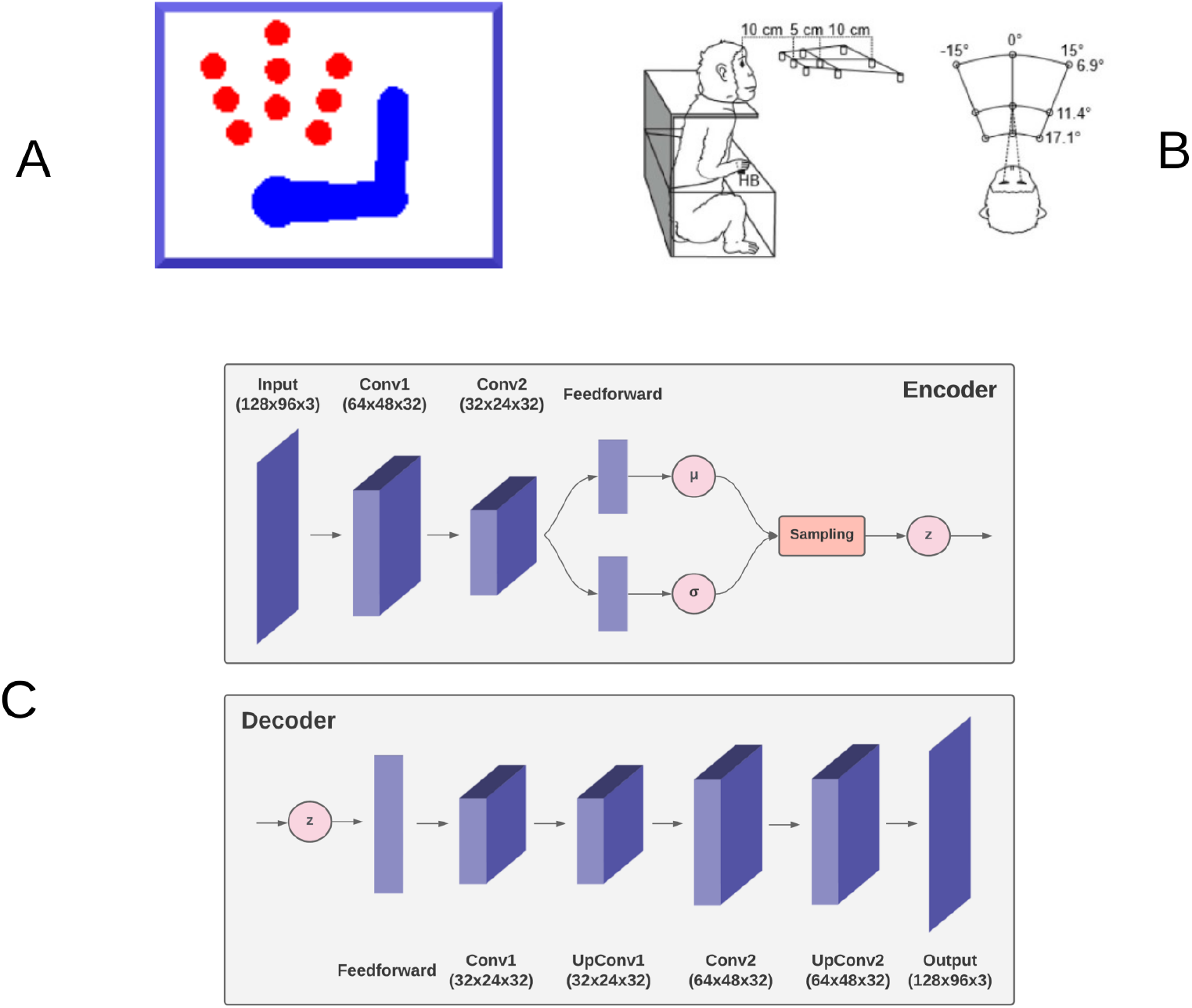
Simulation outline. The agent, a simulated 3-DoF actuated upper limb shown in (A) is set to reach one of the nine red circle targets as in the reference monkey experiment [58] outlined in (B). The agent is equipped with a fixed virtual camera providing peripersonal visual input and a visual model, the decoder ***g**_v_* of a VAE shown in (C) simulating functions of the DVS.

In the following, we first describe the agent and the specific implementation. Then, we assess of the agent’s perceptual and motor control capabilities in static and dynamic conditions. The static condition simulated a typical monkey reaching task of peripersonal targets as in Figure 3 [58]. In turn, the dynamic condition involved a moving target that the agent had to track continuously.

#### 3.4.1 Body

The body consists of a simulated monkey upper limb composed of a moving torso attached to an anchored neck, an upper arm, and a lower arm, as shown in Figure 3. The three moving segments are schematized as rectangles, each with unit mass, while the joints (shoulder, elbow) and the tips (neck, hand) as circles. The proportions of the limb segment and the operating range of the joint angles were derived from monkey data *Macaca mulatta* [67]. The state of the limb and its dynamics are described by the joint angles ***θ*** and their first moment 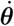. We assume noisy velocity-level motor control, whereby the motor efferents ***a*** noisily control the first moment of joint angles with zero-centered Gaussian noise:

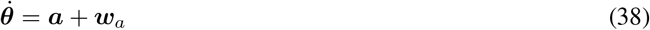

#### 3.4.2 Sensors

The agent receives information about its proprioceptive state and visual context. Simplified peripersonal visual input ***s**_v_* was provided by a virtual camera that included three 2D color planes, each 128×96 pixels in size. The location and orientation of the camera were fixed so that the input provided full vision of peripersonal targets and the entire limb in any possible limb state within its operating range. The limb could occlude the target in some configurations.

As in the simulated limb, the motor control system also receives proprioceptive feedback through sensors ***s**_p_,* providing noisy information on the true state of the limb [68, 49]. We further assumed that ***s***_p_ provides a noisy reading of the state of all joints only in terms of joint angles, ignoring other proprioceptive signals such as force and stretch [24], which the Active Inference framework can natively incorporate.

#### 3.4.3 Belief

We assume that the belief ***μ*** comprises three components: (i) beliefs ***μ**_a_* over arm joint angles, or posture; (ii) beliefs ***μ**_t_* over the target location represented again in the joint angles space - i.e., the posture corresponding to the arm touching the target; and (iii) beliefs ***μ**_h_* over a memorized home button (HB) position. Thus, ***μ*** = [***μ**_a_, **μ**_t_, **μ**_h_*]. Note that the last two components can be interpreted as *affordances*, allowing the agent to implement interactions in terms of bodily configurations [69].

#### 3.4.4 Intentions

Stepping on the proposed formalization (Equation 20), we define two specific intentions (Figure 4) as follows:

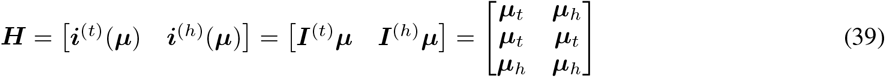

**Figure 4:**
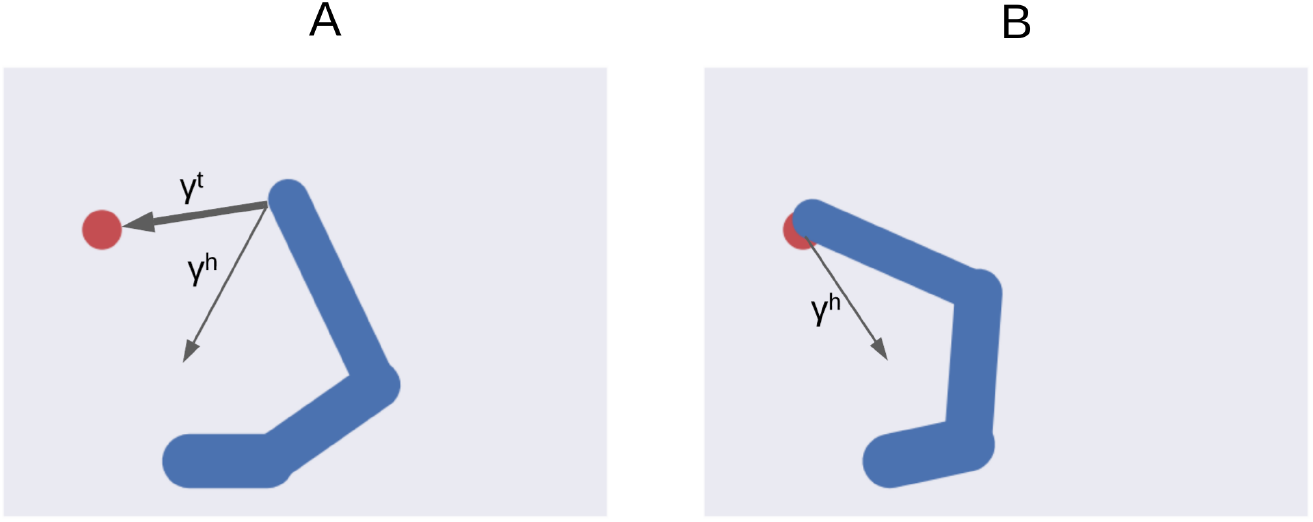
Multiple intentions. Intention prediction errors of variable strength (arrow width) controlled by intention precisions point at different target states. (A) A stronger target attraction (the red circle) implements the reaching action. (B) A stronger attraction of the invisible but previously memorized HB implements the return-to-home action.

Here ***h***^(*t*)^ = (***μ***) defines the agent’s expectation that the arm belief is equal to the joint configuration corresponding to the target to be reached, and it is implemented as a simple mapping ***I***^(*t*)^ that sets the first belief component equal to the second one. In turn, the intention ***h***^(*h*)^ = ***i***^(*h*)^ (***μ***) encodes the future belief of the agent that the arm will be at the HB position. The two intention mappings are defined by:

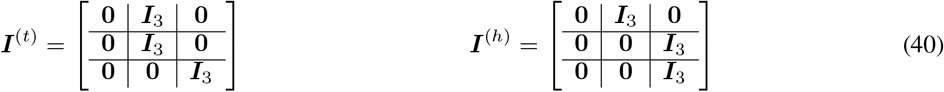

where **0** and ***I***_3_ are respectively 3×3 zero and identity matrices. The corresponding intention prediction errors are then:

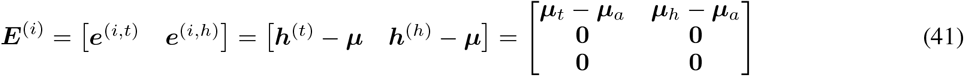

These errors provide an update direction respectively toward the target and HB joint angles. As there is no intention to move the target or the HB, the second and third components of the prediction errors will be zero.

#### 3.4.5 Sensory Predictions

The sensory generative distribution has two components, one for each sensory modality: a simplified proprioceptive model ***g**_p_*(***μ***) and a full-blown visual model ***g**_v_* (***μ***):

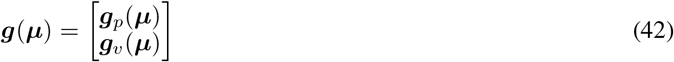

Since the belief is already in the joint angles domain, we implemented a simple proprioceptive generative model ***g**_p_*(***μ***) = ***G**_p_**μ*** = ***μ**_a_*, where ***G**_p_* is a mapping that only extracts the first component of the belief:

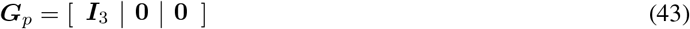

Note that ***g**_p_* (***μ***) could be easily extended to a more complex proprioceptive mapping if the body and/or joint sensors have a more complex structure and the belief has a richer and abstract representation.

In turn, the visual generative model ***g**_v_* is the decoder component of a VAE (see Figure 3C) that produces a visual representation of both target and body configurations given the overall belief in joint angles (example in Figure 13). The decoder consists of one feedforward layer, two transposed convolutional layers, and two standard convolutional layers needed to smooth the output.

The gradient **G** of the full generative model is the following:

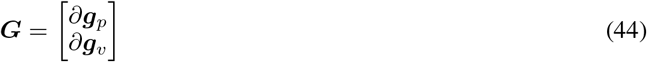

Here, the proprioceptive gradient simplifies to the proprioceptive mapping **G**_p_ itself, while the visual component is the gradient of the decoder computed by backpropagation.

Therefore, predictions ***p*** and prediction errors ***E**_s_* take the form:

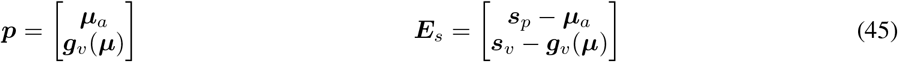

Note that defining the sensory predictions on both proprioceptive and visual sensory domains allows the agent to perform efficient goal-directed behavior also in conditions of visual uncertainty, e.g., due to low visibility. Indeed, since the belief is maintained over time, the agent remembers the last known target position and can thus accomplish reaching tasks also in case of temporarily occluded targets.

#### 3.4.6 Precisions

Free energy minimization and Predictive Coding in general heavily depend on precisions. To investigate their role, we parameterized the relative precisions of each intention and sensory domain with parameters *α* and *β* as follows:

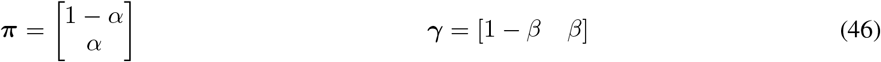

The parameter *α* controls the relative strength of the error update due to proprioception and vision, while the parameter *β* controls the relative attraction by each intention. With these parameters, the sensory and intention prediction errors are unpacked as follows:

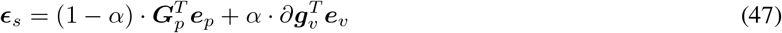

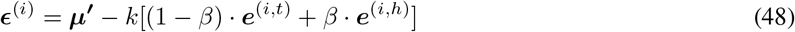

Equation (47) shows the balance between visual and proprioceptive information. For example, if *α* = 0, the agent will only use proprioceptive feedback, while if *α* =1, the belief will be updated only relying on visual feedback. Note that these are extreme conditions - e.g., the former may correspond to null visibility - and typical sensory systems provide balanced feedback.

In turn, Equation (48) spells out the control of belief attraction. The agent will follow the first intention when *β* = 0, or the second one when *β* = 1 (Figure 4). Intermediate intention strengths will lead to an attractor conflict, thus we assume that biased competitive inhibition implements the control of intention selection. Finally, the parameter k controls the overall attractor magnitude (see also Equation 26).

We can also use the precision parameter α to manipulate the strength of the free energy derivative with respect to the actions as follows:

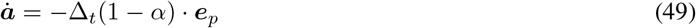

Note that by increasing *α* - i.e., more reliability on vision - the magnitude of belief update remains constant, while action updates decrease because the agent becomes less confident about its proprioceptive information. Also, one could differentially investigate the effect of precision strength on belief and action by directly manipulating specific precisions; e.g., visual precisions ***π**_v_* may include different components that follow the belief structure:

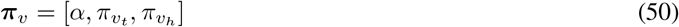

For example, when *α* = 0 and *π_v_t__* > 0, the target belief is updated using visual input while the arm moves only using proprioception, a scenario that emulates movement in darkness with a lit target.

### 3.5 Computational Analysis

In the following, we assess the capacities of the intention-driven Active Inference agent to perceive and perform goal-directed actions in reaching tasks with static and dynamic visual targets. The main testbed task was delayed reaching, but we simulated several other conditions.

Sensorimotor control that implements goal-directed behavior was investigated in various sensory feedback conditions, including pure proprioceptive or mixed visual and proprioceptive, in which the VAE decoder provided support for dynamic estimation of visual targets and bodily states. The latter is the typical condition of performing reaching actions and allows greater accuracy [70]. In an additional *baseline* (BL) condition, the target was estimated by the decoder, but the movement was performed without visual feedback or proprioceptive noise, to allow comparisons with the typical approach in previous continuous Active Inference studies, e.g., [53]. We also investigated the effects of sensory and intention precisions, motor control type, and movement onset policy. Finally, we analyzed the visual model and the nature of its gradients to provide critical information about the causes of the observed motor behavior.

Action performance was assessed with the help of several measures: (i) *reach accuracy:* success in approaching the target within 10 pixels of its center, i.e., the hand touching the target; (ii) *reach error: L*^2^ hand-target distance at the end of the trial; (iii) *reach stability:* standard deviation of *L*^2^ hand-target distance during the period from target reach to the end of the trial, in successful trials; (iv) *reach time*: number of time steps needed to reach the target in successful trials. We also assessed target perception through analog measures based on the *L*^2^ distance between the target location and its estimate transformed from joint angles into visual position by applying the geometric (forward) model. Specifically, we defined the following measures: (v) *perception accuracy:* success in estimating the target location within 10 pixels; (vi) *perception error: *L*^2^* distance between the true and estimated target position at the end of the trial; (vii) *perception stability*: standard deviation of the *L*^2^ distance between the target position and its estimation during the period starting from successful estimation until end of the trial; (viii) *perception time:* number of time steps needed to successfully estimate the target position.

#### 3.5.1 Delayed Reaching Task

The primary testbed task is a simplified version of a delayed reaching monkey task in which a static target must be reached with a movement that can only start after a delay period [58]. Delayed actions are used to separately investigate neural processes related to action preparation (e.g., perception and planning) and execution in goal-directed behavior, and are thus useful to analyze the two main computational components of free energy minimization, namely, perceptual and active inference, which otherwise work in parallel. Delayed reaching could be implemented using various approaches: the update of the posture component of the belief dynamics could be blocked by setting the intention gain *k* to 0 during inference (implemented here): in this way, there are no active intentions and the belief only follows sensory information. Alternatively, action execution could be temporarily suspended by setting the parameter *α* to 1, so that the agent still produces proprioceptive predictions but does not trust the corresponding prediction errors: in this scenario, the belief dynamics includes a small component directed toward the intention, but the discrepancy produced is not minimized through movement.

Reach trials start with the hand placed on the HB located in front of the body center (i.e., the “neck”), and the belief is initialized with this configuration. Then one of the nine possible targets of the reference experiment (Figure 3) is lit red. Follows a delay period of 100 time steps during which the agent is only allowed to perceive the visible target and the limb, and the inference process can only change the belief. After that, the limb is allowed to move and the joint angles are updated according to (38). As in the reference task, upon target reaching the agent stops for a sufficiently long period, i.e., a total of 300 time steps per trial. After that, the agent reaches back the HB (not analyzed here). The simulation included 100 repetitions per target, i.e., 900 trials in total.

As described before, we assume that the HB position is encoded and memorized as a belief during previous experiences, and therefore used to compute the (competing) reaching intention. However, this creates a conflict among intentions aiming to fulfill opposing goals, while the agent can physically realize only one of them at a time (Figure 4). We therefore set the parameter *β* (Equation 48) to realistic values of 0.1 to activate the reaching intention and 0.9 to implement the HB return intention. Such parameter may be however dynamically modulated through higher-level bias and mutual inhibition.

Figure 5 illustrates key points of the delayed reaching task. During the delay period (A-C), the posture does not change since the joint angles only follow the arm belief, which is kept fixed, while the target belief is attracted by the sensory evidence and gradually shifts toward it. When movement is allowed (D-F) by setting k > 0 and β = 0.1, the combined attractor produces a force that moves the arm belief toward the target, generating proprioceptive predictions - therefore motor commands - that let the real arm follow such trajectory.

**Figure 5:**
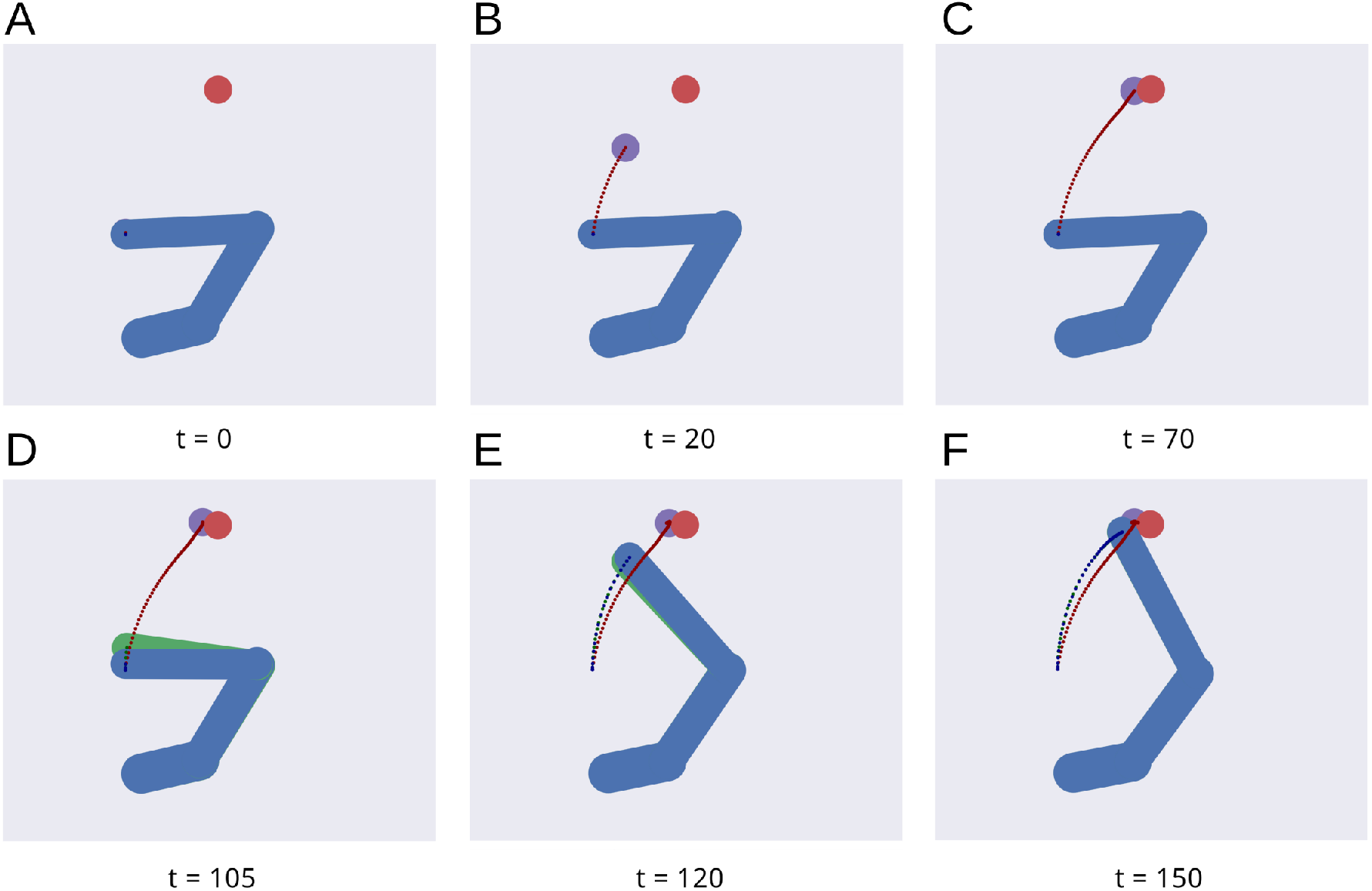
Dynamics of the delayed reaching task. At trial onset (A) a visual target (red circle) appears, the arm (in blue) is located on the HB position, and the arm belief (in green) is set at the true arm state. During the delay period, the perceptual inference process gradually drives the target belief (purple circle) toward the real position (B, C). During this phase, the intention gain k is set to 0, so that movement is inhibited and the arm belief does not change given the unchanged proprioceptive evidence. After movement onset, the arm freely follows its belief (D, E) until they both arrive at the goal state (F).

Reaching performance is summarized in Figure 6. Panels A-D show spatial statistics of the final hand location (with the corresponding belief) for each target, separately for reaching with proprioception only or proprioceptive and visual sensory feedback. Descriptive statistics revealed an important benefit of visual feedback (E-H), in parallel with classical behavioral observations [70]: reach accuracy was higher (with: 88.28%; without: 83.72%) and both reach stability and arm belief error were considerably better with visual feedback as well (stability: 1.35; error: 1.98px) compared to the condition with only proprioception (stability: 1.78; error: 2.87px).

**Figure 6:**
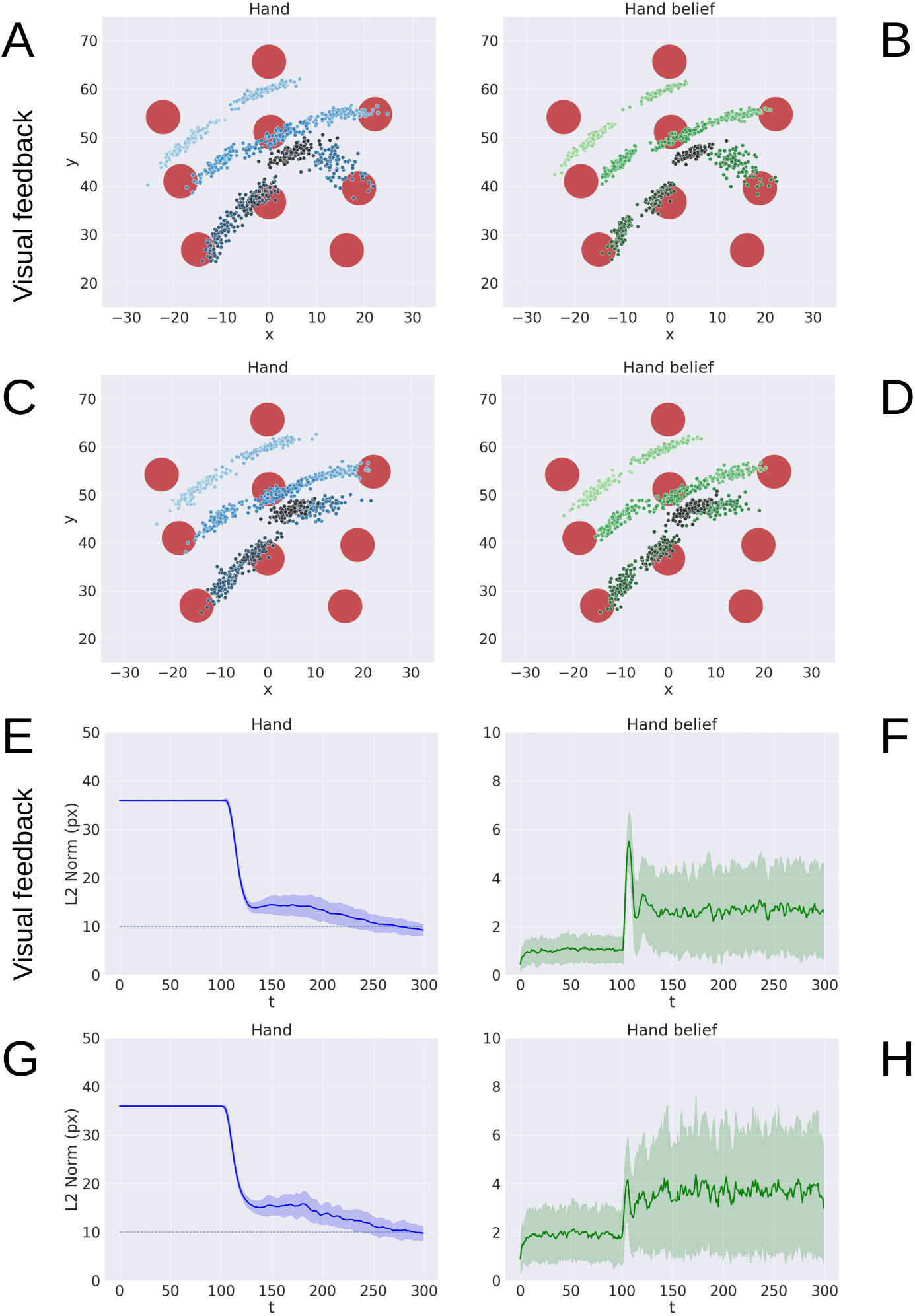
Performance of the delayed reaching task. (A-D) Spatial distribution of hand positions (A, C) and corresponding beliefs (B, D) per target at the end of the reach movements, with (A, B) and without (C, D) visual feedback. Each point represents a trial (100 trials per target). Reach error (E, G) and belief error (F, H) over time, with (E, F) and without (G, H) visual feedback (bands represent C.I.). The reach criterion of the hand-target distance is visualized as a dotted line. *L*^2^ norm for the hand belief is computed by the difference between real and estimated hand positions. Reaching with visual feedback resulted in a more stable hand belief.

#### 3.5.2 Precision Balance

The effects of sensory feedback led to a further systematic assessment of the effects of sensory and intention precisions, denoted by *α, π_v_t__* and k (see Equations 46 and 50). The assessment was carried out following the structure of the delayed reaching task. We varied the above precisions one at a time, using levels shown on the abscissas in Figure 7, while keeping the non-varied precisions at their default values. Note that *α* = 0 corresponds to reaching without visual feedback, while the conditions *α* > 0 may be interpreted as reaching with different levels of arm visibility. We recall that the baseline condition (BL) performs reaching movements without visual feedback and proprioceptive noise, i.e., *α* = 0 and *w_p_* = 0. To obtain a systematic evaluation, each condition was run on a rich set of 1000 randomly selected targets that covered the entire operational space. Finally, we only considered the target-reaching intention, i.e., β = 0; everything else was the same as in the main task.

**Figure 7:**
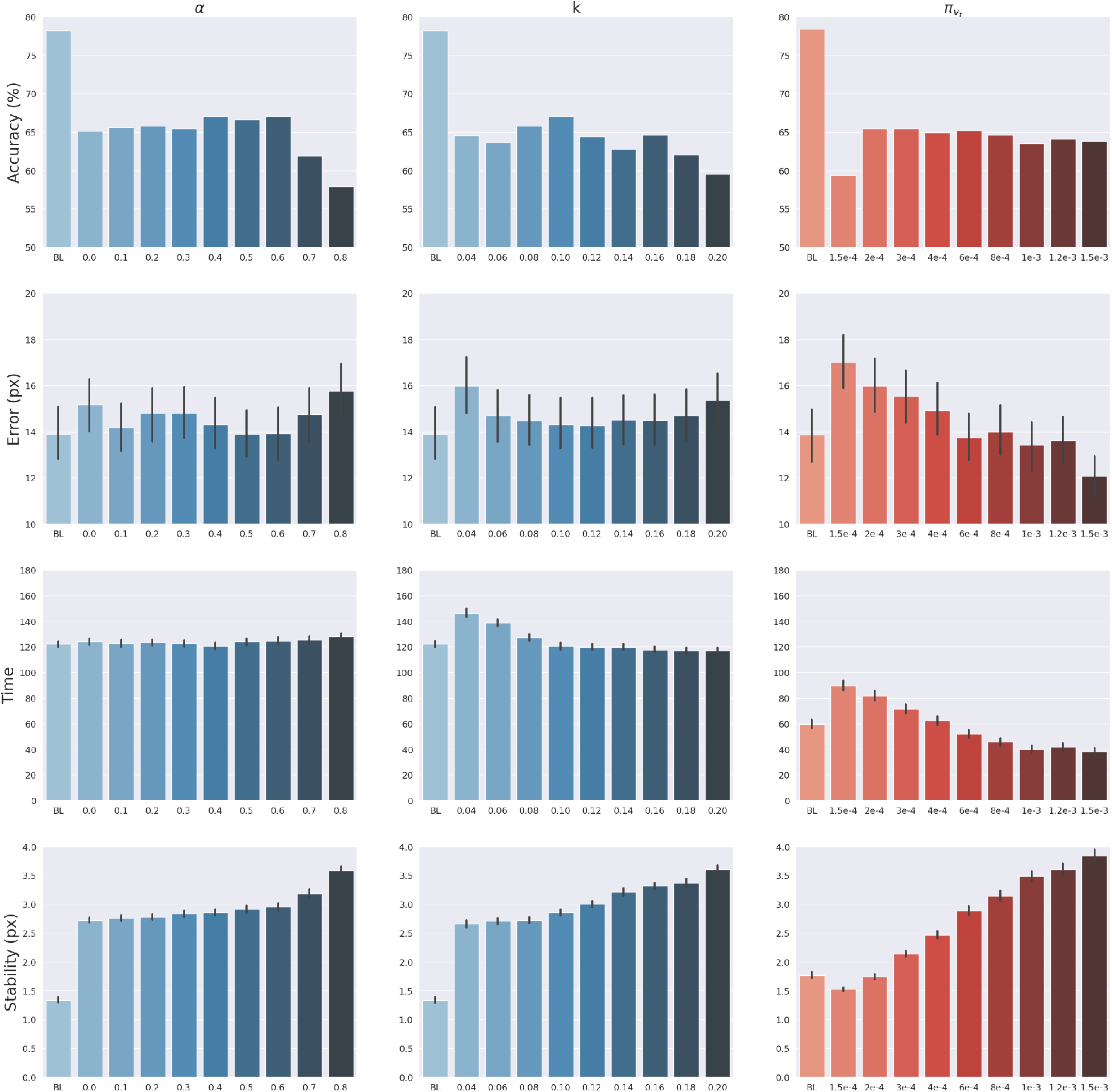
Effects of precision balance. Reach accuracy (1st row), reach error, i.e., *L*^2^ hand-target distance (2nd row), reach time (3rd row), and reach stability (4th row). Reach performance is shown as a function of arm sensory precision *α* (left) and attractor strength *k* (middle). In turn, perception performance is shown as a function of target sensory precision *π_v_t__* (right). Vertical bars represent C.I.

The results are shown in Figure 7. The panels in the left column show the effect of *α* compared to the BL agent with noiseless proprioception. Active Inference with only proprioception (i.e., *α* = 0) has a lower reach accuracy and higher error, while the best performance is obtained with balanced proprioceptive and visual input, in corroboration with the observations of the basic delayed reaching task. In the latter case, the motor control circuitry continuously integrates all available sensory sources to implement visually-guided behavior [71]. However, accuracy and stability rapidly decrease for excessively high values of a, due to the discrepancy in update directions between the belief - which makes use of all available sensory information, including the more precise visual feedback - and action - which in this case relies on excessively noisy proprioception.

Furthermore, as in the main experiment, the effects of visual precision are evident in the stability of the arm belief, which gradually improves with increasing values (Figure 8): In addition to the reliability of the visual input, this effect is also a consequence of the smaller action updates due to the reduced proprioceptive precision.

**Figure 8:**
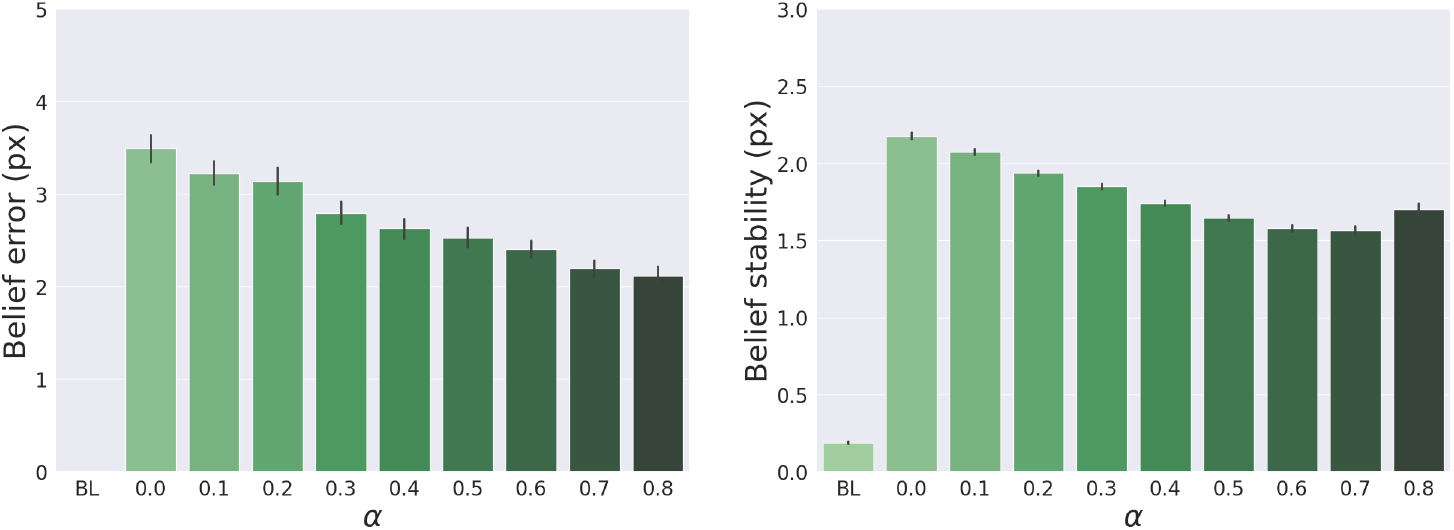
Belief error and stability - representing the difference between real and estimated hand positions - for different values of *α*. Vertical bars represent C.I.

In turn, the panels in the middle column of Figure 7 reveal the effects of the attractor gain k; to remind the reader, the greater the gain, the greater the contribution of intention prediction errors to belief updates. The results show that as the intention gain k increases reach accuracy generally improves, and the number of time steps needed to reach the target decreases. However, beyond a certain level, the accuracy tends to decrease since the trajectory dynamics becomes unstable; thus, excessively strong action drag is counterproductive to the implementation of smooth movements.

Finally, the panels in the right column of Figure 7 show the effects of the target precision *π_v_t__*, which directly affects the quality of target perception. Note that better performances are generally obtained in terms of accuracy, error, and perception time for values of *π_v_t__* higher than the arm visual precision, which corresponds to a classical effect of contrast on perception, but also means here that the arm and target beliefs follow different dynamics.

#### 3.5.3 Motor Control

We described earlier two different ways of implementing motor control in Active Inference: making use of all sensory information, or proprioception only. The first method requires significantly more computations since the agent needs to know the inverse mapping from every sensory domain to compute the control signals. However, given the assumptions we made, this approach is potentially more stable because it updates both belief and action with the same information. On the other hand, a pure proprioception-based control mechanism could produce potentially incorrect movements because the motor control commands result from comparing proprioceptive predictions with noisy observations. Greater cost-effectiveness of the second method thus might come at the cost of worsened performances, which we investigate here.

Figure 9 shows a comparison of the two control methods and the BL agent, evaluated under the same conditions we used to investigate precision balance, including 1000 random targets. Performance was measured in terms of belief and reach stability and reach accuracy. The results reveal, first, that the expected decreased belief stability of the full model with respect to the BL agent (C) does not affect hand stability (B), although the proprioceptive noise apparently contributed to decreased reach accuracy (A). More importantly, the results confirm our expectations that pure proprioception control has considerably lower reach stability caused by incorrect update directions of the motor control signals, resulting in a greater decreased reach accuracy relative to the full model.

**Figure 9:**
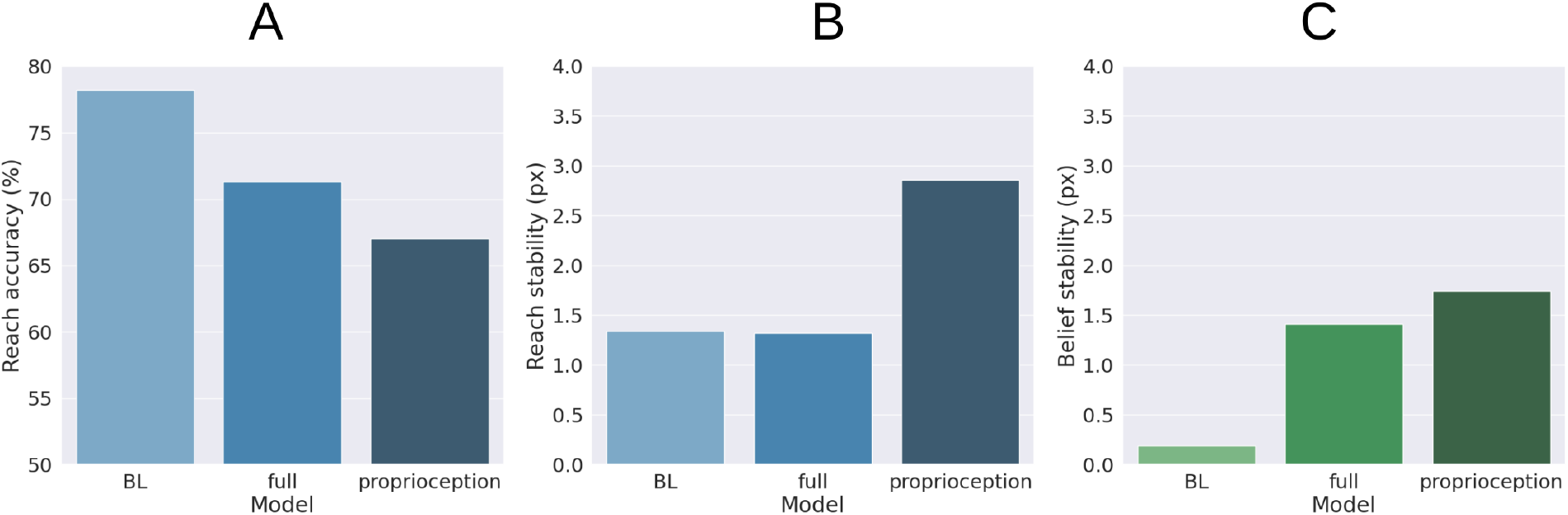
Motor control methods. Reach accuracy (A), reach stability (B), and belief stability (C) for a BL agent and the two different implementations of motor control, based either on all sensory information (full control) or on proprioception only.

#### 3.5.4 Movement Onset Policy

We also investigated the effects of movement onset using several policies, which differ by the duration of the period of pure perception preceding full Active Inference. One such policy we investigate here is characteristic of actions performed under time pressure, in which movement starts along with perception, i.e., action is *immediate*. Another policy that could be considered typical for acting under normal conditions has movement beginning with the satisfaction of a certain perception criterion. This policy *dynamically* deliberates the onset of movement. Various perception criteria could be used: here, the action starts when the norm of the target belief 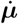*_t_* remained below a given threshold (i.e., 0.01) for a certain period (i.e., 5 time steps). These parameters were arbitrarily chosen in consideration of exploratory delayed reaching simulations. Finally, we include the previously used delayed action policy in which movement onset is delayed by a *fixed* period (here, 100 time steps, sufficient to obtain a precise target estimation). To obtain systematic observations, each policy was again run on 1000 randomly selected targets. Measurements included reach and perception accuracy, motor control stability after reach, target perception stability, as well as reach time since the beginning of the trial or after movement onset.

Figure 10 shows the results with the three different policies. Although the reach error is approximately the same in all tested conditions, the agent controlled by the immediate policy reached the target within the lowest total number of time steps: target perception and intention setting were dynamically computed along with movement onset. However, if we consider the total task time, the number of time steps is the highest in this condition, since the arm belief and the arm itself move along with the slow visual target estimation.

**Figure 10:**
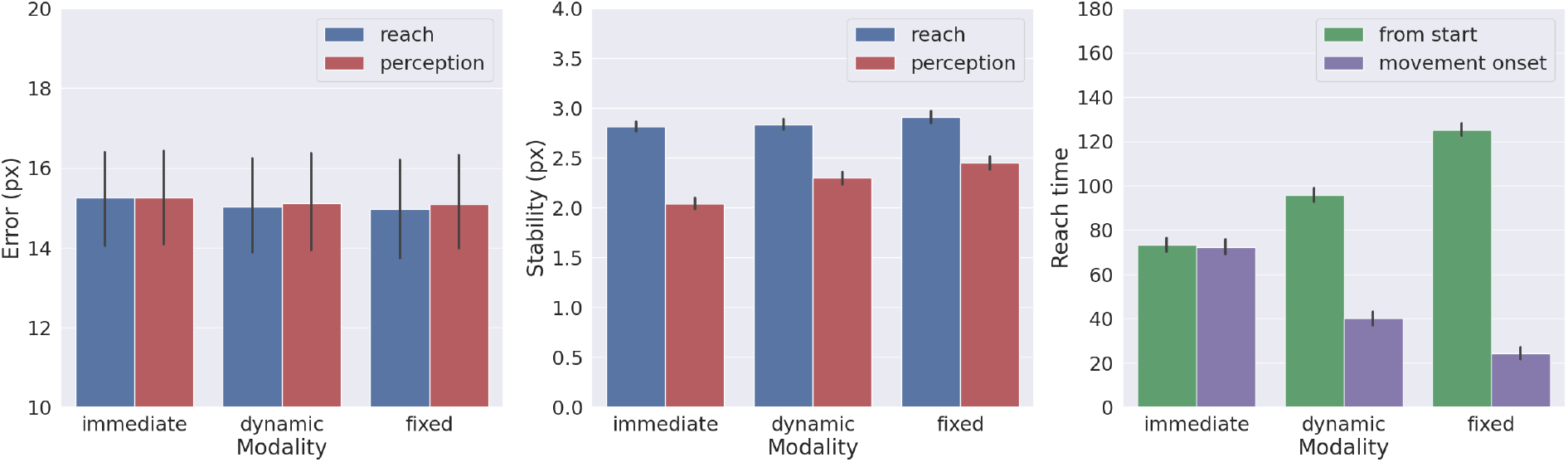
Effects of movement onset policy. Reach error (left), stability (middle), and time (right) across several policies (immediate, dynamic, and fixed delay). Vertical bars represent C.I.

In turn, if the agent starts the movement when the uncertainty about the target position is already minimized (either in the *dynamic* or *fixed* condition), the movement time decreases, although if added on top of the perception time results in slower actions relative to the immediate movement condition. Finally, we note that target perceptual stability somewhat decreases for dynamic and fixed policies; this somewhat unexpected result is encouraging for dynamic target tracking tasks in which immediate movement onset is mandatory.

#### 3.5.5 Tracking Dynamic Targets

In a second testbed task, the agent was required to track a smooth-moving target whose initial location was randomly chosen from the entire operational space. In each trial, the targets received an initial velocity of 0.1px per step in a direction uniformly spanning the 0-360 degree range. When the target reached a border, its movement was reflected. As in the previous simulations, the belief was initialized at the HB configuration and the trial time limit was 300 time steps. However, for the agent to correctly follow the targets, both the belief and action were dynamically and continuously inferred in parallel, i.e., without a pure perceptual period.

Figure 11 shows the reach trajectory in dynamic target tracking for 10 random trials. The left panel shows the evolution over time of *L*^2^ hand-target distance, while the right panel represents the error between estimated and true target positions. The results suggest that the agent is generally able to correctly and dynamically estimate the beliefs over both target and arm for almost every trial, also in the case of moving targets. In some cases however, mainly when the target is out of reach, it is temporarily or permanently “lost” in terms of its belief, which has also the consequences of losing the target in terms of reach. Further analysis with a more realistic bodily configuration and visual sensory system - as well as comparisons with actual kinematic data - should provide further insights into the capabilities of Active Inference to perform dynamic reaching.

**Figure 11:**
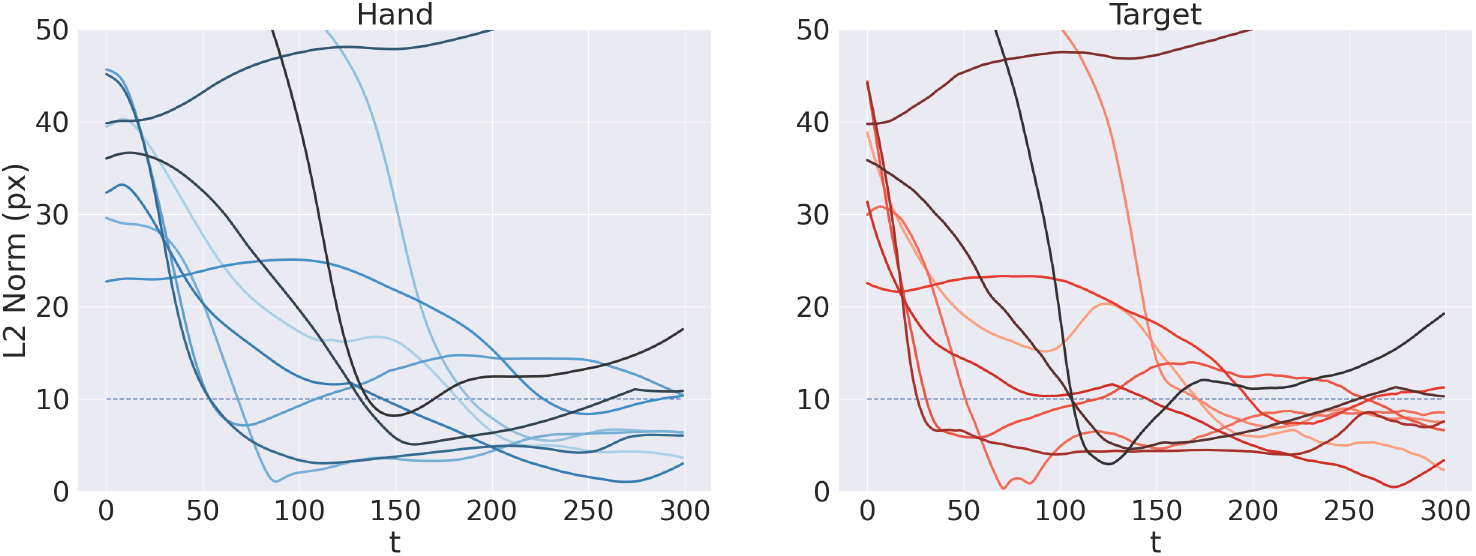
Tracking dynamic targets. Reach (left) and perception (right) error over time for 10 random trials.

#### 3.5.6 Free Energy Minimization

Here we illustrate the dynamics of free energy minimization in delayed reaching, which is at the heart of continuous Active Inference. To that aim, we run 10 new reaching trials with static and dynamic targets and recorded the free energy derivatives with respect to generalized belief and action.

Figure 12A-F shows the trajectory of the free energy derivatives with respect to the arm and target components during delayed reaching of a static target; the two columns show the trends for the last two joints, i.e., the arm and forearm segments, that most strongly articulate the reaching action.

**Figure 12:**
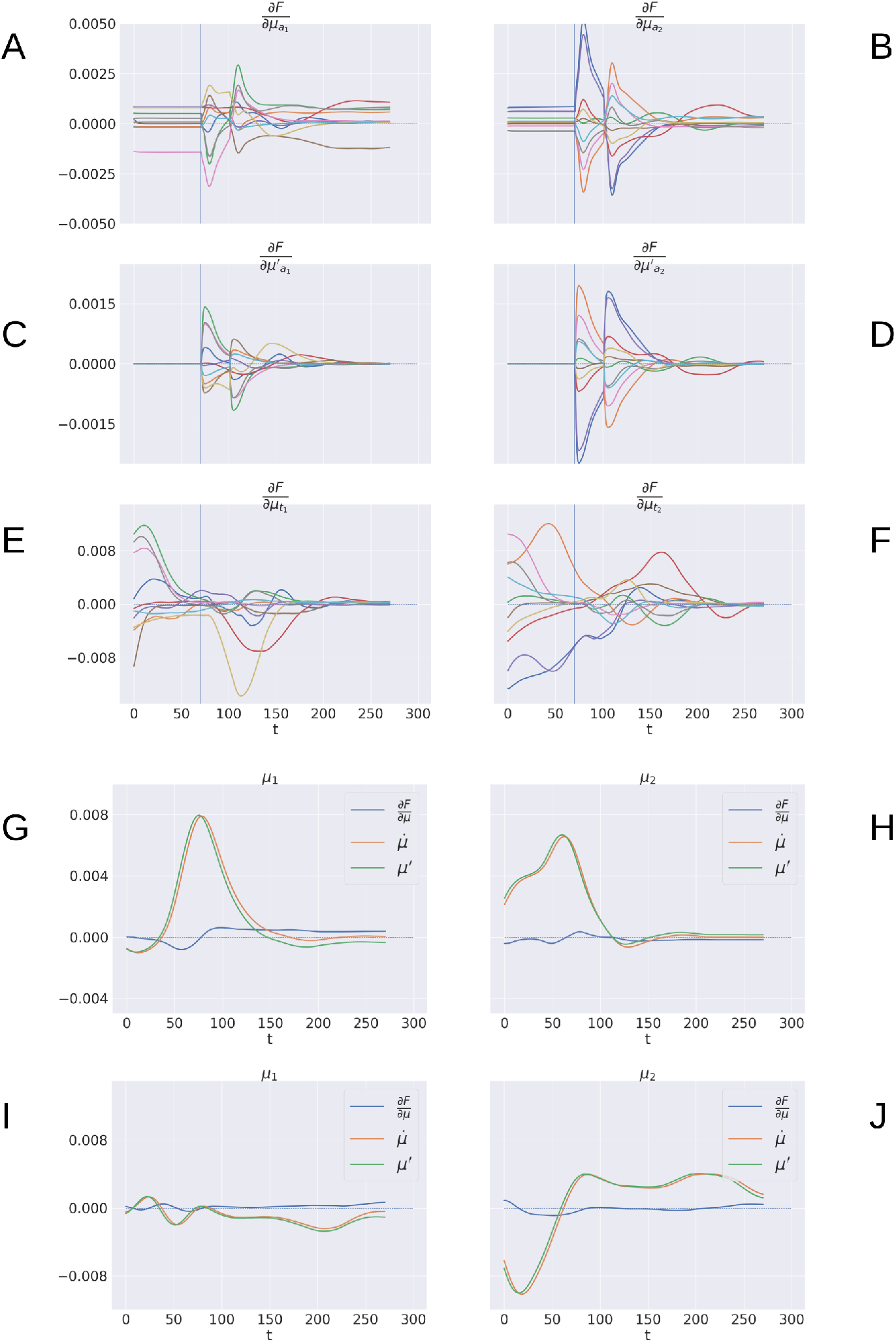
Free energy minimization. (A-F) Free energy derivative with respect to the 0th- and 1st-order belief for arm (A-D) and target (E-F). (G-J) Comparison between the reference frames of the belief - the path of the mode 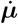 and the mode of the path ***μ’*** - for sample static (G-H) and dynamic (I-J) trials. The left/right columns refer to the arm/forearm segments. Trials data are smoothed with a 30 time-step moving average.

Note that the gradients of the free energy with respect to the target belief are rapidly minimized during the initial perceptual phase while the arm gradients remain still. Upon action execution (indicated by a vertical line), the arm gradients rapidly change as well, resulting in updated proprioceptive predictions that drive arm movements. However, arm movements causes change on the visuals scene, which have a secondary effect over the just-minimized free energy on target belief.

Figure 12G-J goes even deeper, showing a direct comparison between 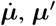, ***μ’***, and the difference 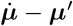, on sample static (G-H) and dynamic (I-J) targets. We recall that free energy minimization implies that the two reference frames (the path of the mode 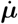 and the mode of the path ***μ’***) should overlap at some point in time, when the agent has inferred the correct trajectory of the generalized hidden states. This is crucial especially in dynamic reaching, in which the aim is to capture the instantaneous trajectory of every object in the scene. The decreasing free energy gradients (blue lines) show that this aim is indeed successfully achieved in both static and dynamic tasks.

#### 3.5.7 Visual Model Analysis

Here, we provide an assessment of the visual model whose performance is critical for accurate visually-guided motor control. To recall, the visual model is implemented with a VAE trained offline to reconstruct images of arm-target configurations such as the one in Figure 13A. A critical VAE parameter is the variance of the recognition (encoder) density Σ***_ϕ_*** (see Equation 17). We therefore evaluated its effect on perception and action by training several VAEs with different variance levels. VAEs were trained for 100 epochs on a dataset comprising 20.000 randomly drawn body-target configurations that uniformly spanned the entire operational space. The target size varied with a radius ranging from 5 to 12 pixels. VAE performance was assessed on other 10.000 randomly selected configurations that uniformly sampled the space, with a target size of 5 pixels (the default condition for the Active Inference tests).

**Figure 13:**
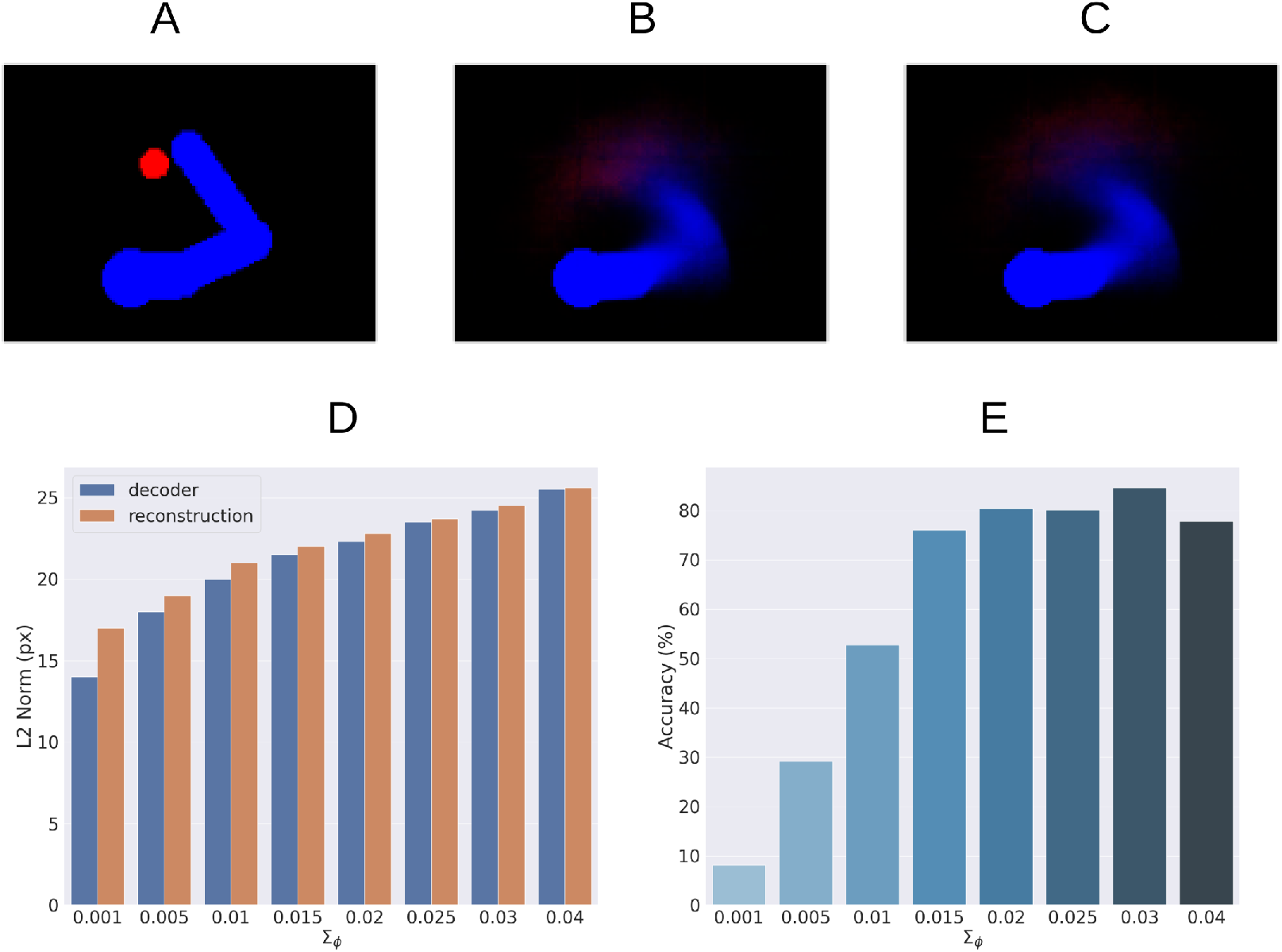
Visual model analysis. (A-C) Sample visual observation (A) and its decoding from joint angles (B) and through complete encoding-decoding process (C). (D-E) Visual model performance. Quality of perception measured as the *L*^2^ norm between observed and reconstructed images (D) and accuracy of Active Inference (E), as a function of the recognition density variance *Σ_ϕ_*.

Most critical was the VAE capacity to generate adequate images of joint arm-target configurations, which we measured with the help of the *L*^2^ norm between visual observations, and VAE-generated images. To provide more insights on the two VAE processes, decoding and encoding, we proceeded as follows: first, *decoding* was assessed by generating images for given body-target states such as that in Figure 13B. The decoded images were compared with the ground truth images produced by applying the geometric model for the same state of the body target (Figure 13A). Second, full VAE performance was assessed by computing the average *L*^2^ norm between observed images and their full VAE *reconstruction*, i.e., first encoding and then decoding them (as in Figure 13C). Third, we directly assessed the specific effects of the recognition density variance on Active Inference using the BL condition of the delayed reaching task as a measure.

Figure 13D represents the results of the perceptual assessment tests, showing the *L*^2^ norm between the original and generated images as a function of recognition density variance. As expected, lower variances generally resulted in lower errors with respect to both pure decoding and full encoding-decoding. Surprisingly, however, the accuracy of Active Inference in the reaching task behaved somewhat differently: the best accuracy was obtained not for predictions with low variance, but for intermediate variance levels (Figure 13E). This could be explained by the fact that low-variance images imply highly non-linear gradients that prevent correct gradient descent on free energy. On the other hand, as the variance increases the reconstructed image becomes somewhat blurred, which helps obtain a smoother gradient that correctly drives free energy minimization and therefore improves movement accuracy (more on this in the next section). However, as the variance continues to increase, the reconstructed images become too blurry, degrading both belief inference and motor control.

#### 3.5.8 Visual Gradient Analysis

To further investigate the cause of the unexpected low variance issue, we analyzed the consistency of the visual gradient *∂**g**_v_* of the decoder for several encoder variance values. To this aim, we computed the gradients for different reference states over the entire operational space.

Figure 14A-C reveals that a decoder with intermediate variance values (green line) causes smaller but smoother gradients, while a too-low variance (orange line) causes sharp peaks near the reference point and even incorrect gradient directions in some cases. Therefore, too low encoder variances seem to make the decoder prone to overfitting, while higher variance values help extract a smoother relationship between irregular multidimensional sensory domains and regular low-dimension causes.

**Figure 14:**
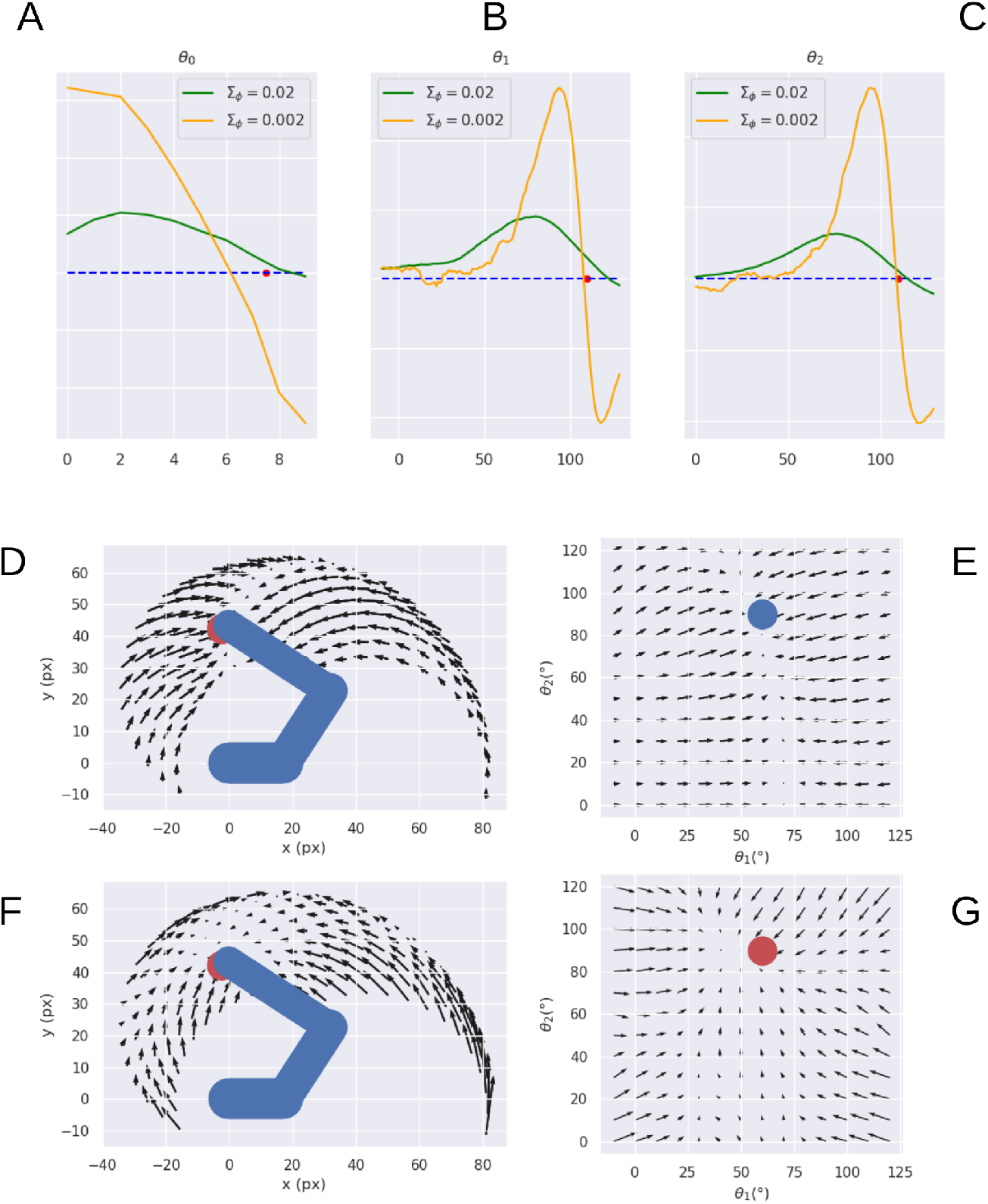
Visual gradient analysis. Marginal gradients for each joint, i.e., neck (A), shoulder (B), and elbow (C), and for two values of the recognition density variance Σ*_ϕ_* (green/orange line) computed by backpropagating the error between images with different arm configurations (abscissa: joint angle) and a reference image (whose angle is represented by the red dot on the abscissa). (D-G) Gradients for arm (D, E) and target (F, G) in both Cartesian (D, F) and joint (E, G) space.

Figure 14D-G further illustrates the arm and target gradients relative to a sample reference posture and target location (the result is similar for other configurations) in both Cartesian and polar coordinates; the polar plot shows the two joints most relevant to the reaching action.

The plots reveal greater arm gradients (upper panels) in the vicinity of the target location; in that subspace, the decoder has less uncertainty about which direction to choose to minimize the error. Notably, the gradients tend to compose curved directions, a characteristic of biological motions. The polar plot provides critical insights into the causes of the circular pattern: the gradients are mostly parallel to the horizontal axis, which corresponds to a movement consisting essentially of pure shoulder rotation. Thus, they provide a strong driving force on the shoulder almost throughout the operational space, while the area in which the elbow is controlled is limited to the vicinity of the target location. These gradients result in a two-phase reaching of static targets in which the agent first rotates the shoulder - resulting in a horizontal positioning - and then starts to rotate the elbow as soon as the latter enters its attraction area. The same gradients can explain the linear motion pattern of an arm tracking dynamically moving targets when the arm is close to the target: in that case, all gradients provide motion force as explained above.

On the other hand, the gradients of the target belief (bottom panels) behave somewhat differently: since this belief is unconstrained and can freely move in the environment, update directions more directly approach the target in all angular coordinates (see the polar plot to the right). Yet, linear belief updates in the polar space still translate to curve directions in the Cartesian space.

## 4 Discussion

We presented a normative computational theory based on Active Inference of how neural circuitry in the PPC and DVS supports visually-guided actions in a dynamically changing environment. Our focus was on the computational basis of encoding dynamic action goals in the PPC through flexible motor intentions. The theory is based on Predictive Coding [38, 37] and Active Inference [72] and evidence suggesting that the PPC performs visuomotor transformations [10, 11, 12] and encodes motor plans [3, 4]. Accordingly, the PPC is proposed to maintain dynamic expectations of both current and desired latent states over the environment and use them to generate proprioceptive predictions that ultimately generate movements through reflex arcs [36, 49]. In turn, the DVS encodes a generative model that translates latent state expectations into visual predictions. Discrepancies between sensory-level predictions and actual sensations produce prediction errors sent back through the cortical hierarchy to improve the internal representation. The theory unifies research on intention coding [4] and current views that the PPC estimates the body and environmental states [18], providing specific computational hypotheses regarding the involvement in goal-directed behavior. It also extends some perception-bound Predictive Coding interpretations of the PPC dynamics [73] and provides a more comprehensive account of movement planning [1], tightly integrated into the overall sensorimotor control process.

How does the proposed model compare with state-of-the-art implementations of continuous Active Inference? First, we designed an internal belief over not only bodily states but also every object in the scene. The latter beliefs are encoded in the joint angles space as well, simulating a visuomotor reference frame that the PPC is supposed to encode. Second, we decomposed the belief dynamics into a set of independent intentions depending on the current belief and predicting the next plausible state. Such formalization has several advantages: (i) as attractors are dynamically generated at each time step, the agent can also follow moving targets and interact with a constantly changing environment; (ii) letting the future goal state depend on the current belief at the same hierarchical level allows to change only specific components, while keeping the others fixed to their current values or free to change by prediction errors coming from other directions of the cortical hierarchy; (iii) expressing the target position in terms of a possible joint configuration - either imposed by higher levels for realizing specific object affordances or freely inferred by the exteroceptive models - results in a simple intention, without the need of directly using sensory information or duplicating lower-level generative models, which results in implausible scenarios; (iv) maintaining different belief components also allows easy encoding of previously memorized states which can be especially useful when implementing a sequence of actions, since only the intention precisions have to be adjusted. Indeed, it seems that the PPC explicitly encodes and maintains such goals during the whole unfolding of sequential actions [63].

A specific goal is selected among other competitive intentions possibly under the control of the PFC and PMd [65] and fulfilled by setting it as a predominant belief trajectory as an attractor with a strong gain (see Equations 41 and 26 and Figure 4). For example, in a typical reaching task, the goal of reaching a specific visual target corresponds to the future expectation that the agent’s arm will be over that target; thus, if the agent maintains a belief over it, the corresponding intention links the expected belief over the future body posture with the inferred target, expressed in joint angles, encoding a specific interaction to realize.

In light of these theoretical considerations, the theory predicts several different types of correlates in the PPC related to environment, task, and body states. The former two include correlates of potential spatial targets and selected motor goals, which indeed have been consistently found in the PPC [3, 4, 15]. The latter includes correlates of intention-biased bodily state estimates which thus are not precise representations of the true states. Furthermore, the use of generalized beliefs in Active Inference predicts that the PPC encodes not only static states but also a detailed estimate of body dynamics, up to a few temporal orders. Indeed, a body of literature report motion-sensitive, or Vision-for-Action activity in the DVS and PPC [11]. A last set of correlations regards prediction errors. In fact, body-environment transitions involving a change of states and tasks result in transient bursts of activity in error-conveying cells until the error is minimized. Prediction errors conveying upstream information are putatively encoded by pyramidal cortical cells in superficial layers while downstream predictions are encoded by deep pyramidal neurons [74].

We tested the computational feasibility of the theory on a delayed reaching task - a classical experiment in electrophysiology - in which a monkey is required to reach with its hand a visual-spatial target, starting the movement from an HB [58]. To do this, we simulated an agent consisting of a coarse 3-DoF limb model and noisy visual and proprioceptive sensors (Figure 3A). Simplified proprioceptive sensors provided a noisy reading of the state of the limb in joint angles, while visual input was provided by a fixed camera and consisted of an image of the target and limb. Predictive visual sensory processing simulating the DVS was implemented with a VAE trained to infer body state and target location, both in the joint angles domain (Figure 3C). The limbs were animated at the velocity level with motor control signals computed by the visually-guided Active Inference controller.

The computational analysis showed, first and most importantly, that the controller could correctly infer the position of the visual targets (Figure 5, *t*=70), use it to compute and set motor goals in terms of prior beliefs on the future body state through intention functions (Figure 5, *t*=105), and perform adequate and smooth reach movements (Figure 5 *t*=105-150), with and without visual feedback (Figure 6). The greater accuracy obtained with visual feedback parallels classical results in a similar classical behavioral comparison of reaching [70].

We then systematically investigated the effects of noise on various functional components (Figure 7), starting with the balance of the precision between proprioceptive and visual sensory models: a noiseless Active Inference agent (BL condition) resulted in the best performance, with a stable final approach and accuracy only limited by the quality of the visual target estimation. Among the noisy conditions, pure proprioceptive control resulted in the lowest performance, as expected. Motor control driven by both proprioceptive and visual feedback with balanced precision between the two domains resulted in improved reach accuracy and greatly improved arm belief stability (Figure 8). The effect on accuracy was mainly due to the inclusion of visual information in the inference process, but also to slower updates of the motor control signals due to decreased confidence about proprioceptive input.

The increased stability of the arm belief did not improve movement stability as increasing confidence about visual input also increased the discrepancy between belief and action updates, the latter only relying on noisy proprioceptive observations. In fact, we showed that if we remove the plausibility constraint that motor control is driven only by proprioceptive predictions and thus let actions minimize prediction errors from all sensory domains, the reach performance greatly increases (Figure 9). Nonetheless, any combination of visual and proprioceptive feedback improved performance relative to a control driven by feedback from a single sensory domain.

The instability due to the difference in update directions between belief and action could be balanced by other mechanisms that we have not considered here. For example, we assumed that the same pathway is used for both control and belief inference, but it seems that the motor cortex generates different predictions depending on the brain areas which it interacts with: purely proprioceptive predictions for motor control, whose prediction error is suppressed at the lowest level of the hierarchy, and rich somatosensory predictions for latent state inference, which integrates somatic sensations at different hierarchical levels [36].

Intention precisions or attractor gains affected performance as well. First, they affected reach time: as expected, the greater the gain, the faster the movement. However, fast movements come at a cost: increased gains generally resulted in less precise movements and decreased stability during the final reach period. Finally, higher visual target precisions decreased perception time and improved perception accuracy but decreased perception stability.

We also investigated the effects of movement onset policies: response delay allows investigating perceptual and motor preparatory processes separately from the motor control and action execution. We found that delayed response decreased movement time with respect to a policy that requires an immediate response (Figure 10), which fits the behavioral pattern [75]. Apparently, this is due to the need of estimating, in the latter condition, the target position “on the fly”, and constantly adapting the intention according to the updated target estimate. The advantage of allowing some preparatory time becomes clear in an anecdotal fly-catching task, which results in faster movement and increased chances of success. This comes with the critical contribution of PPC neurons that systematically modulate their activity during the preparatory period [75], which here provided specific predictions for the computations performed in the PPC.

Notably, the immediate-response policy allowed the Active Inference controller to perform actions under dynamic environmental conditions, such as tracking moving objects. Free energy minimization resulted in rapid target detection also in this case, and maintained subthreshold perception error on moving targets (Figure 11) which allowed precise tracking after an initial reaching period.

Intention-driven Active Inference in continuous time largely compares to classical neural-level hypotheses of motor planning such as the Dynamic Field Theory [1], with the advantage of stepping on an established Predictive Coding framework, dynamic approximate probabilistic inference, and end-to-end sensorimotor control. The Dynamic Field Theory estimates the parameters of the desired movement - such as movement direction and target velocity - from sensory and task features encoding environmental descriptors, which closely compares to motor goal coding through flexible intentions in our model. The two theories have in common a dynamic activation of the internal representations in continuous time, governed by a dynamic system, but they differ in the nature of the signals and their coding. Movement descriptors in Dynamic Field Theory are represented by a dynamically activated multidimensional space, each encoded on a population of competing neurons, while Active Inference approximates movement properties with their central moments (belief) and dispersion (precisions). While population coding allows a complete description of a probabilistic distribution, it could be overshot when used to code single magnitudes (although it is essential when encoding discrete categories). Yet, such representation allows coding multiple competing targets on the same population of neurons, while in our scheme each target should be encoded by a dedicated unit. Notably, the brain encodes scalar variables using a variety of number coding schemes, including monotonic and distributed [76]. The latter, known as a “mental number line” [77], could be an interesting hypothesis to explore also in the context of feature coding in continuous Active Inference. Currently, distributed coding is used only in discrete Active Inference and other probabilistic models to investigate computationally high-level cognitive functions such as planning, navigation, and control [29, 65, 78]. The two theories also differ in the nature of the input to their dynamic systems. In Active Inference, system input encodes generalized prediction errors, which are integrated into higher-level moments. Instead, input in Dynamic Field Theory directly encodes state values. Coding based on prediction errors has the advantage of minimizing the quantity of transmitted information - hence, energy. Finally, the theories also differ in scope: Active Inference provides a full account of the entire sensorimotor control process, while the Dynamic Field Theory describes only movement planning.

### 4.1 Precision Balance and Conditions for Disorders

Based on our computational analysis, it becomes clear that some motor and behavioral disorders could be due to the lack of proper sensory and intention precisions [79]. Here, we illustrate the normal condition and two types of potentially improper precision balance that could become a causal condition for neurological disorders.

Figure 15A illustrates the condition for normal functioning, which is such that the contribution of a single intention to the belief update (which, as a reminder to the reader, is proportional to the gain of intentions k) is sufficiently small with respect to the sensory contribution. In this case, during free energy minimization, the system dynamics smoothly moves the belief toward the strongest goal, along with precise tracking of the true latent state and sensory signal of the limbs, allowing thus to compute correct motor control errors and perform smooth action execution.

**Figure 15:**
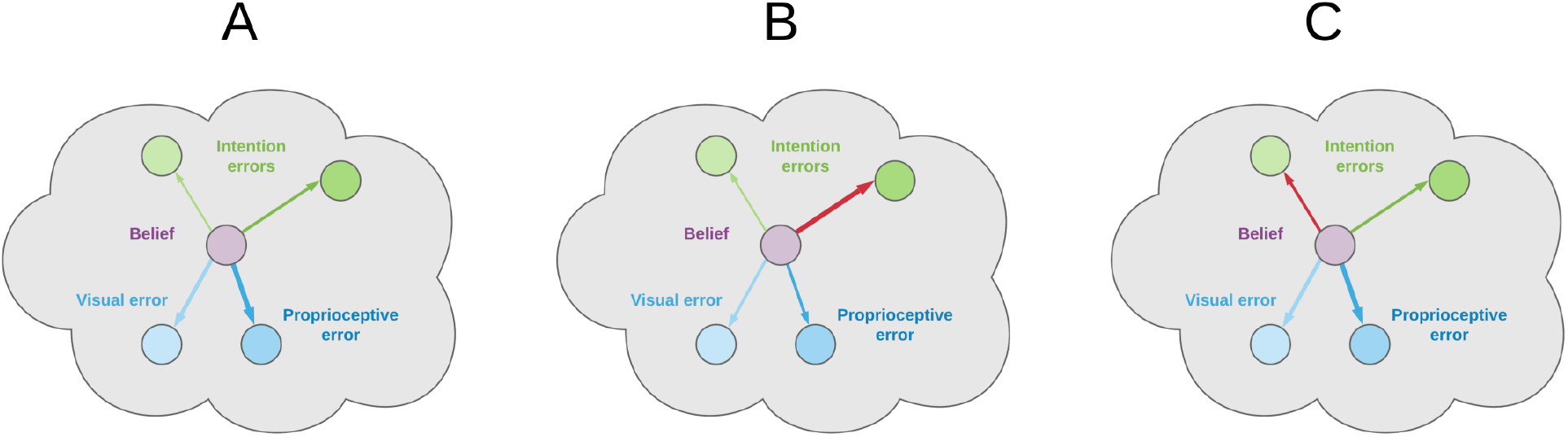
Normal and abnormal intention gains and precisions. Arrow widths represent strengths of prediction errors. In normal conditions, the sensory contribution to free energy minimization is bigger than that of the intentions and there is a clear unambiguous intention to fulfill (A). An abnormally strong intention gain k drives the belief away from the true joint states (B). Too close precision levels ***γ*** also hinder implementing competing intentions (C).

A critical abnormal condition arises when the intention gain k is too strong, as illustrated in Figure 15B. In this case, the belief moves too rapidly toward the goal without being able to match the proprioceptive observations, which results in computing incorrect motor control signals. Another abnormal condition is caused by too close precisions **γ** of competitive intentions, which is likely to result in opposing belief updates and thus prevent the fulfillment of any of the competing goals (as in Figure 15C). This situation might manifest in terms of motor onset failure or oscillatory behavior.

### 4.2 Limitations and Future Directions

Our focus here was on intention coding in the PPC, which directly deals with motor plans and motor control. Further elaborations will extend the theory with higher-level aspects of cognitive control, including intention structuring (dealt by PM), phasing (SMA) [7], planning, and goal selection (HC, PFC) [65, 80, 81].

Motor control operated here in an inner belief space belonging to the joint angles domain, which is generally suboptimal in the external Cartesian space. In fact, we assumed that neural activity in the PPC encodes generalized beliefs over targets and body only in a motor-related domain; however, neural data suggest that neurons in the motor cortex encode motor trajectories also in extrinsic coordinates [36, 14], and a more realistic model should include representations encoding states in both intrinsic and extrinsic reference frames. A functional correlate of the motor cortex should represent future states - which were defined here implicitly in the intention prediction errors and dynamics functions - and transform desired trajectories from Cartesian coordinates to proprioceptive predictions in the intrinsic state-space. This transformation is different from Optimal Control planning since the optimization of a classical inverse model reduces to a more manageable inference problem. Finally, although free energy minimization in a single belief domain is faster and more parsimonious, more precise visually-guided motor control should include continuous activation of visual low-level attractors at a certain “neural energy” cost since motor areas interact with and generate predictions in multiple sensory domains. The current work just partially covers the three main domains of sensorimotor learning through a predictive forward control; for example, it does not fully include reactive, stimuli-driven control such as obstacle avoidance, although we showed that it can successfully perform static and dynamic tasks.

Since our focus was on the theoretical introduction of intentionality in Active Inference, every analysis was only partially characterized by a simple reaching task. However, fundamental properties of the physical model, including geometry, mass, and friction, strongly influence the resulting motion dynamics - hence the entire inferential process. This implementation does not adopt other important neural and biomechanical specificities such as signal delay and joint friction [82]. However, it could be easily extended to accommodate additional sensory modalities - e.g., tactile sensations - with rich generative models such as the VAE here implemented. Further planned computational analyses will use a richer belief space, a more realistic physical arm model, and additional actuators, and expand the complexity of the intention functions to investigate the capacity of the theory to explain in-depth neural levels, cognitive, and kinematic phenomena related to motor learning, motion perception, motor planning, and so on.

Planned future studies with a more articulated agent will also challenge the theory at the behavioral and neural level against other empirical findings regarding movement preparation and motor control, in either delayed or direct response settings. For example, we will test the model for stimulus-stimulus congruency and stimulus-response compatibility effects [83]. As for the former, it is intuitive that a greater sensory dimension overlap predicts faster target-belief convergence - thus faster intention setting. Less intuitive is that stimulus-response compatibility effects should emerge due to differences in the dynamic transition from one belief state to another in the proprioceptive domain. For example, a belief over the effector state should change more, requiring more time to converge when reaching a contralateral position than an ipsilateral one.

Although we considered an Active Inference model with just a single layer of intentions, the structure represented in Figure 2 could be scaled hierarchically and consider intermediate goals between high-level intentions and low-level sensory generative models, e.g., by combining discrete and continuous Active Inference for planning and movement execution [84, 85, 86, 87]. According to the free energy principle, the agent will then choose goals and subgoals and rely on specific sensory modalities such that free energy is minimized at every hierarchical level based on prediction errors coming from the level below. This formalization will provide an explicit basis for motor planning, including tasks like object manipulation. Indeed, although the current implementation performs well on spatial tasks like reaching in a dynamically changing environment, it cannot implement composite goals which the brain needs to handle. On the other hand, an agent that can encode higher-level intentions in a discrete domain and infer policies based on the *expected free energy* will be able to dynamically modify its behavior and react to environmental changes. An extended implementation of this kind will be an interesting and more biologically plausible alternative to standard reinforcement learning algorithms.

## Acknowledgments

This research received funding from the European Union’s Horizon 2020 Framework Programme for Research and Innovation under H2020-EIC-FETPROACT-2019 Grant Agreement 951910 (MAIA) to IS, Grant Agreement No 945539 (Human Brain Project SGA3), the European Research Council under Grant Agreement No. 820213 (ThinkAhead), and from the Italian Ministry for Research MIUR under Grant Agreement PRIN 2017KZNZLN (PACE) to IS.

## Notes

### Competing Interest Statement

The authors have declared no competing interest.

### Summary of Updates

In this revision we discuss more background literature and further clarify the significance of the result. We also put more emphasis on the computational aspect of the result by slightly changing the title and the claims about the putative neural substrate.

## References

[1] Wolfram Erlhagen and Gregor Schöner. Dynamic field theory of movement preparation. Psychological Review, 109(3):545–572, 2002.

[2] Rajesh P.N. Rao and Dana H. Ballard. Predictive coding in the visual cortex: A functional interpretation of some extra-classical receptive-field effects. Nature Neuroscience, 2(1):79–87, 1999.

[3] Richard A. Andersen. Encoding of intention and spatial location in the posterior parietal cortex. Cerebral Cortex, 5(5):457–469, 1995.

[4] L. H. Snyder, A. P. Batista, and R. A. Andersen. Coding of intention in the posterior parietal cortex. Nature, 386(6621):167–170, 1997.

[5] Leonardo Fogassi, Pier Francesco Ferrari, Benno Gesierich, Stefano Rozzi, Fabian Chersi, and Giacomo Rizzolotti. Parietal lobe: From action organization to intention understanding. Science, 308(5722):662–667, 2005.

[6] Hakwan C Lau, Robert D. Rogers, Patrick Haggard, and Richard E. Passingham. Attention to Intention. Sicence, 303(5661):1208–1210, 2004.

[7] Juan A. Gallego, Tamar R. Makin, and Samuel D. McDougle. Going beyond primary motor cortex to improve brain–computer interfaces. Trends in Neurosciences, 45(3):176–183, 2022.

[8] L. H. Snyder, A. P. Batista, and R. A. Andersen. Intention-related activity in the posterior parietal cortex: A review. Vision Research, 40(10-12):1433–1441, 2000.

[9] Karl J Friston, Jérémie Mattout, and James Kilner. Action understanding and active inference. Biological cybernetics, 104(1-2):137–60, feb 2011.

[10] Patrizia Fattori, Rossella Breveglieri, Annalisa Bosco, Michela Gamberini, and Claudio Galletti. Vision for Prehension in the Medial Parietal Cortex. Cerebral Cortex, 27(February):1149–1163, 2017.

[11] Claudio Galletti and Patrizia Fattori. The dorsal visual stream revisited: Stable circuits or dynamic pathways? Cortex, 98:203–217, 2018.

[12] Paul Cisek and John F Kalaska. Neural Mechanisms for Interacting with a World Full of Action Choices. Annual Review of Neuroscience, 33:269–298, 2010.

[13] Michela Gamberini, Lauretta Passarelli, Matteo Filippini, Patrizia Fattori, and Claudio Galletti. Vision for action: thalamic and cortical inputs to the macaque superior parietal lobule. Brain Structure and Function, 226(9):2951–2966, 2021.

[14] Yale E. Cohen and Richard A. Andersen. A common reference frame for movement plans in the posterior parietal cortex. Nature Reviews Neuroscience, 3(7):553–562, 2002.

[15] Matteo Filippini, Rossella Breveglieri, Kostas Hadjidimitrakis, Annalisa Bosco, and Patrizia Fattori. Prediction of Reach Goals in Depth and Direction from the Parietal Cortex. Cell Reports, 23(3):725–732, 2018.

[16] M Desmurget, C M Epstein, R S Turner, C Prablanc, G E Alexander, and S T Grafton. PPC and visually directing reaching to targets. Nature Ne, 2(6):563–567, 1999.

[17] Claudio Galletti, Michela Gamberini, and Patrizia Fattori. The posterior parietal area V6A: An attentionally-modulated visuomotor region involved in the control of reach-to-grasp action. Neuroscience and Biobehavioral Reviews, 141(August):104823, 2022.

[18] W. Pieter Medendorp and Tobias Heed. State estimation in posterior parietal cortex: Distinct poles of environmental and bodily states. Progress in Neurobiology, 183(April):101691, 2019.

[19] Shlomi Haar and Opher Donchin. A revised computational neuroanatomy for motor control. Journal of Cognitive Neuroscience, 32(10):1823–1836, 2020.

[20] Maurizio Corbetta and Gordon L. Shulman. Control of goal-directed and stimulus-driven attention in the brain. Nature Reviews Neuroscience, 3(3):201–215, 2002.

[21] Reza Shadmehr and John W. Krakauer. A computational neuroanatomy for motor control. Experimental Brain Research, 185(3):359–381, 2008.

[22] Emanuel Todorov. Optimality principles in sensorimotor control. Nature Neuroscience, 7:907–915, 2004.

[23] Meel Velliste, Sagi Perel, M. Chance Spalding, Andrew S. Whitford, and Andrew B. Schwartz. Cortical control of a improsthetic arm for self-feeding. Nature, 453(7198):1098–1101, 2008.

[24] Shriya S Srinivasan, Samantha Gutierrez-Arango, Ashley Chia En Teng, Erica Israel, Hyungeun Song, Zachary Keith Bailey, Matthew J. Carty, Lisa E. Freed, and Hugh M. Herr. Neural interfacing architecture enables enhanced motor control and residual limb functionality postamputation. Proceedings of the National Academy of Sciences of the United States of America, 118(9), 2021.

[25] Karl Friston and Stefan Kiebel. Predictive coding under the free-energy principle. Philosophical Transactions of the Royal Society B: Biological Sciences, 364(1521):1211–1221, 2009.

[26] Karl J. Friston, Jean Daunizeau, James Kilner, and Stefan J. Kiebel. Action and behavior: A free-energy formulation. Biological Cybernetics, 102(3):227–260, 2010.

[27] Rafal Bogacz. A tutorial on the free-energy framework for modelling perception and learning. Journal of Mathematical Psychology, 76:198–211, 2017.

[28] Thomas Parr, Giovanni Pezzulo, and Karl J Friston. Active inference: the free energy principle in mind, brain, and behavior. Cambridge, MA: MIT Press, 2021.

[29] Giovanni Pezzulo, Francesco Rigoli, and Karl J. Friston. Hierarchical Active Inference: A Theory of Motivated Control. Trends in Cognitive Sciences, 22(4):294–306, 2018.

[30] Karl J Friston, S Samothrakis, and Read Montague. Active inference and agency: optimal control without cost functions. Biological cybernetics, (106):523–541, 2012.

[31] Raphael Kaplan and Karl J. Friston. Planning and navigation as active inference. Biological Cybernetics, 112(4):323–343, 2018.

[32] Marc Toussaint and Amos Storkey. Probabilistic inference for solving discrete and continuous state Markov Decision Processes. ACM International Conference Proceeding Series, 148:945–952, 2006.

[33] Beren Millidge, Alexander Tschantz, Anil K. Seth, and Christopher L. Buckley. On the relationship between active inference and control as inference. Communications in Computer and Information Science, 1326:3–11, 2020.

[34] Sergey Levine. Reinforcement Learning and Control as Probabilistic Inference: Tutorial and Review. 2018.

[35] Karl Friston. What is optimal about motor control? Neuron, 72(3):488–498, 2011.

[36] Rick A. Adams, Stewart Shipp, and Karl J. Friston. Predictions not commands: Active inference in the motor system. Brain Structure and Function, 218(3):611–643, 2013.

[37] Jakob Hohwy. The Predictive Mind. Oxford University Press UK, 2013.

[38] Kenji Doya. Bayesian Brain: Probabilistic Approaches to Neural Coding. 2007.

[39] Giovanni Pezzulo, Francesco Donnarumma, Pierpaolo Iodice, Domenico Maisto, and Ivilin Stoianov. Model-based approaches to active perception and control. Entropy, 19(6), 2017.

[40] Christopher M Bishop. Pattern Recognition & Machine Learning. Springer, New York, New York, USA, 2006.

[41] Wei Ji Ma, Jeffrey M Beck, Peter E Latham, and Alexandre Pouget. Bayesian inference with probabilistic population codes. Nature Neuroscience, 9(11):1432–1438, 2006.

[42] Karl Friston, Jérémie Mattout, Nelson Trujillo-Barreto, John Ashburner, and Will Penny. Variational free energy and the Laplace approximation. NeuroImage, 34(1):220–234, 2007.

[43] Karl Friston. Hierarchical models in the brain. PLoS Computational Biology, 4(11), 2008.

[44] K. J. Friston, N. Trujillo-Barreto, and J. Daunizeau. DEM: A variational treatment of dynamic systems. NeuroImage, 41(3):849–885, 2008.

[45] Karl Friston. The history of the future of the Bayesian brain. NeuroImage, 62(2):1230–1233, 2012.

[46] Christopher L. Buckley, Chang Sub Kim, Simon McGregor, and Anil K. Seth. The free energy principle for action and perception: A mathematical review. Journal of Mathematical Psychology, 81:55–79, 2017.

[47] Manuel Baltieri and Christopher L. Buckley. PID control as a process of active inference with linear generative models. Entropy, 21(3), 2019.

[48] Thomas Parr and Karl J. Friston. The anatomy of inference: Generative models and brain structure. Frontiers in Computational Neuroscience, 12(November), 2018.

[49] Christopher Versteeg, Joshua M. Rosenow, Sliman J. Bensmaia, and Lee E. Miller. Encoding of limb state by single neurons in the cuneate nucleus of awake monkeys. Journal of Neurophysiology, 126(2):693–706, 2021.

[50] Ian J. Goodfellow, Yoshua Bengio, and Aaron Courville. Deep Learning. MIT Press, Cambridge, MA, USA, 2016. http://www.deeplearningbook.org.

[51] Diederik P. Kingma and Max Welling. Auto-encoding variational bayes. 2nd International Conference on Learning Representations, ICLR 2014 - Conference Track Proceedings, pages 1–14, 2014.

[52] Karl J. Friston, Jean Daunizeau, and Stefan J. Kiebel. Reinforcement learning or active inference? PLoS ONE, 4(7), 2009.

[53] Léo Pio-Lopez, Ange Nizard, Karl Friston, and Giovanni Pezzulo. Active inference and robot control: A case study. Journal of the Royal Society Interface, 13(122), 2016.

[54] Jakub Limanowski and Karl Friston. Active inference under visuo-proprioceptive conflict: Simulation and empirical results. Scientific Reports, 10(1):1–14, 2020.

[55] Pablo Lanillos and Gordon Cheng. Adaptive Robot Body Learning and Estimation Through Predictive Coding. IEEE International Conference on Intelligent Robots and Systems, pages 4083–4090, 2018.

[56] Cansu Sancaktar, Marcel A. J. van Gerven, and Pablo Lanillos. End-to-End Pixel-Based Deep Active Inference for Body Perception and Action. In 2020 Joint IEEE 10th International Conference on Development and Learning and Epigenetic Robotics (ICDL-EpiRob), pages 1–8, 2020.

[57] Thomas Rood, Marcel van Gerven, and Pablo Lanillos. A deep active inference model of the rubber-hand illusion. 2020.

[58] Rossella Breveglieri, Claudio Galletti, Giulia Dal Bò, Kostas Hadjidimitrakis, and Patrizia Fattori. Multiple Aspects of Neural Activity during Reaching Preparation in the Medial Posterior Parietal Area V6A. Journal of cognitive neuroscience, 26(4):879–895, 2014.

[59] Rick A. Adams, Eduardo Aponte, Louise Marshall, and Karl J. Friston. Active inference and oculomotor pursuit: The dynamic causal modelling of eye movements. Journal of Neuroscience Methods, 242(May):1–14, 2015.

[60] Mohamed Baioumy, Paul Duckworth, Bruno Lacerda, and Nick Hawes. Active inference for integrated state-estimation, control, and learning. arXiv, 2020.

[61] Guillermo Oliver, Pablo Lanillos, and Gordon Cheng. Active inference body perception and action for humanoid robots. 2019.

[62] Matteo Filippini, Rossella Breveglieri, M. Ali Akhras, Annalisa Bosco, Eris Chinellato, and Patrizia Fattori. Decoding information for grasping from the macaque dorsomedial visual stream. Journal of Neuroscience, 37(16):4311–4322, 2017.

[63] Daniel Baldauf, He Cui, and Richard A. Andersen. The posterior parietal cortex encodes in parallel both goals for double-reach sequences. Journal of Neuroscience, 28(40):10081–10089, 2008.

[64] Aldo Genovesio, Satoshi Tsujimoto, and Steven P Wise. Encoding goals but not abstract magnitude in the primate prefrontal cortex. Neuron, 74(4):656–62, may 2012.

[65] Ivilin Stoianov, Aldo Genovesio, and Giovanni Pezzulo. Prefrontal goal codes emerge as latent states in probabilistic value learning. Journal of Cognitive Neuroscience, 28(1):140–157, 2016.

[66] Andre M. Bastos, W. Martin Usrey, Rick A. Adams, George R. Mangun, Pascal Fries, and Karl J. Friston. Canonical Microcircuits for Predictive Coding. Neuron, 76(4):695–711, 2012.

[67] Yasuhiro Kikuchi and Yuzuru Hamada. Geometric characters of the radius and tibia in Macaca mulatta and Macaca fascicularis. Primates, 50(2):169–183, 2009.

[68] John C. Tuthill and Eiman Azim. Proprioception. Current Biology, 28(5):R194–R203, 2018.

[69] Giovanni Pezzulo and Paul Cisek. Navigating the Affordance Landscape: Feedback Control as a Process Model of Behavior and Cognition. Trends in Cognitive Sciences, 20(6):414–424, 2016.

[70] Steven W. Keele and Michael I. Posner. Processing of Visual Feedback in Rapid Movements. Journal of Experimental Psychology, 77(1):155–158, 1968.

[71] Jeffrey A. Saunders and David C. Knill. Humans use continuous visual feedback from the hand to control fast reaching movements. Experimental Brain Research, 152(3):341–352, 2003.

[72] Karl Friston. The free-energy principle: A unified brain theory? Nature Reviews Neuroscience, 11(2):127–138, 2010.

[73] Thomas H.B. FitzGerald, Rosalyn J. Moran, Karl J. Friston, and Raymond J. Dolan. Precision and neuronal dynamics in the human posterior parietal cortex during evidence accumulation. NeuroImage, 107:219–228, 2015.

[74] Thomas Parr, Giovanni Pezzulo, and Karl J. Friston. Active Inference: The Free Energy Principle in Mind, Brain, and Behavior. The MIT Press, 03 2022.

[75] Krishna V. Shenoy, Maneesh Sahani, and Mark M. Churchland. Cortical Control of Arm Movements: A Dynamical Systems Perspective. Annual Review of Neuroscience, 36(1):337–359, 2013.

[76] Ivilin Stoianov and Marco Zorzi. Emergence of a ‘visual number sense’ in hierarchical generative models. Nature neuroscience, 15(2):194–6, feb 2012.

[77] Ivilin Stoianov, Peter Kramer, Carlo Umiltà, and Marco Zorzi. Visuospatial priming of the mental number line. Cognition, 106(2):770–9, feb 2008.

[78] Ivilin Stoianov, Domenico Maisto, and Giovanni Pezzulo. The hippocampal formation as a hierarchical generative model supporting generative replay and continual learning. Progress in Neurobiology, 217(November 2021):1–20, 2022.

[79] Rick A. Adams, Peter Vincent, David Benrimoh, Karl J. Friston, and Thomas Parr. Everything is connected: Inference and attractors in delusions. Schizophrenia Research, (March), 2021.

[80] I Stoianov, C Pennartz, C Lansink, and G Pezzulo. Model-based spatial navigation in the hippocampus-ventral striatum circuit: a computational analysis. Plos Computational Biology, 14(9):1–28, 2018.

[81] Giovanni Pezzulo, Francesco Donnarumma, Haris Dindo, Alessandro D’Ausilio, Ivana Konvalinka, and Cristiano Castelfranchi. The body talks: Sensorimotor communication and its brain and kinematic signatures. Physics of Life Reviews, 28:1–21, 2019.

[82] Daniel M. Wolpert and J. Randall Flanagan. Computations underlying sensorimotor learning. Current Opinion in Neurobiology, 37:7–11, 2016.

[83] Sylvan Kornblum, Thierry Hasbroucq, and Allen Osman. Dimensional Overlap: Cognitive Basis for Stimulus-Response Compatibility-A Model and Taxonomy. Psychological Review, 97(2):253–270, 1990.

[84] Karl J. Friston, Thomas Parr, and Bert de Vries. The graphical brain: Belief propagation and active inference. 1(4):381–414, 2017.

[85] Karl J. Friston, Richard Rosch, Thomas Parr, Cathy Price, and Howard Bowman. Deep temporal models and active inference. Neuroscience and Biobehavioral Reviews, 77(November 2016):388–402, 2017.

[86] Thomas Parr, Rajeev Vijay Rikhye, Michael M. Halassa, and Karl J. Friston. Prefrontal Computation as Active Inference. Cerebral Cortex, 30(2):682–695, 2020.

[87] Noor Sajid, Philip J. Ball, Thomas Parr, and Karl J. Friston. Active inference: demystified and compared. Neural Computation, 33(3):674–712, 2021.

